# TRAPT: A multi-stage fused deep learning framework for transcriptional regulators prediction via integrating large-scale epigenomic data

**DOI:** 10.1101/2024.05.17.594242

**Authors:** Guorui Zhang, Chao Song, Mingxue Yin, Liyuan Liu, Yuexin Zhang, Ye Li, Jianing Zhang, Maozu Guo, Chunquan Li

**Author notes:** To whom correspondence should be addressed to Chunquan Li.; Maozu Guo. The authors wish it to be known that, in their opinion, the first three authors should be regarded as Joint First Authors.

## Abstract

It is a challenging task to identify functional transcriptional regulators, which control expression of gene sets via regulatory elements and epigenomic signals, involving context-specific studies such as development and diseases. Integrating large-scale multi-omics epigenomic data enables the elucidation of the complex epigenomic control patterns of regulatory elements and regulators. Here, we propose TRAPT, a multi-modality deep learning framework that predicts functional transcriptional regulators from a queried gene set by integrating large-scale multi-omics epigenomic data, including histone modifications, ATAC-seq and TR-ChIP-seq. We design two-stage self-knowledge distillation model to learn nonlinear embedded representation of upstream and downstream regulatory element activity, and merge multi-modality epigenomic features from TR and the queried gene sets for inferring regulator activity. Experimental results on 1072 TR-related datasets demonstrate that TRAPT outperforms current state-of-the-art methods in predicting transcriptional regulators, especially in the prediction of transcription co-factors and chromatin regulators. Additionally, we have successfully identified key transcriptional regulators associated with the disease, genetic variation, cell fate decisions, and tissues. Our method provides an innovative perspective for integrating epigenomic data and has the potential to significantly assist researchers in deepening their understanding of gene expression regulation mechanisms.

## Introduction

The intricate patterns of gene regulation are programmed by multiple upstream transcriptional regulators (TRs), such as transcription factors (TFs), transcription co-factors (TcoFs), and chromatin regulators (CRs), which can mediate regulatory signals between promoters and distal enhancers^1^. The onset of diseases is often associated with aberrant patterns of gene expression, underscoring the importance of identifying the transcriptional regulators that control key gene expression programs. The advancements in ChIP-seq and ATAC-seq techniques have enabled to effectively illustrate cis- and trans-regulatory landscapes. Genomic TRs binding affinities in conjunction with epigenetic information, such as histone modifications and chromatin openness, collectively determine the cell-specific regulatory activities of TRs^2^. Additionally, numerous studies have revealed that transcription factors bind to specific cis-regulatory sequences within the genome, including enhancers and promoters, to modulate the expression of their target genes^.3,4^. Therefore, integrating abundant epigenomic data to identify upstream synergistic regulatory features of genes is imperative for predicting TRs. With the rapid advancement of high-throughput sequencing technologies, a vast accumulation of epigenomic data has been amassed, spanning multiple modalities such as ATAC-seq, DNase-seq, and ChIP-seq, due to differences in sequencing techniques. A major challenge lies in comprehensively collecting and processing these datasets from various sources. Additionally, datasets from different origins exhibit significant issues, including noise interference, batch effects, and data redundancy. Consequently, constructing large-scale data models, developing effective data integration algorithms, and capturing useful data representations while filtering out noise remain substantial challenges.

Some methods have proposed to infer upstream transcriptional regulators using functional gene sets. These include Enrichr^54^, TFEA.ChIP^55^, ChEA3^5^, MAGIC^6^, i-cisTarget^7^, BART^8^, and Lisa^9^. Enrichr, TFEA.ChIP, ChEA3 and MAGIC use gene sets as input, predicting TRs through enrichment analysis with gene sets related to TFs. These approaches essentially involve statistical testing based on the overlaps between the target genes of TRs and the input genes. Although these methods demonstrate notably faster analysis speeds, they do not take into consideration the detailed information of cis-regulatory elements (CREs). As transcription factors function by binding to regulatory elements, information on the cis-regulatory profile is crucial for the accurate inference of regulators. i-cisTarget matches CREs on the genome to predict TF activity through enrichment analysis. i-cisTarget uses CREs instead of merely gene set data, more accurately simulating TF binding, thereby significantly improving the performance of TF activity prediction. However, this algorithm only used CREs associated with input gene set, which is insufficient to simulate the entire genome’s cis-regulatory profile. BART solves the incomplete cis-regulatory profile coverage problem well by inferring the cis-regulatory profile from a large amount of H3K27ac ChIP-seq data using the regression-based MARGE^10^ algorithm. Lisa, known as “MARGE second generation”, further utilizes an extensive array of DNase-seq data, in addition to H3K27ac ChIP-seq data, to enhance the predictive performance of gene-related cis-regulatory profiles. Although BART and Lisa solve the incomplete cis-regulatory profile coverage problem, there is an inherent bias in TR binding, which we refer to as Transcriptional Regulator Binding Preference (TRBP). Essentially, TRs exhibit a predisposition towards associating with regions of active chromatin. More importantly, all existing methods are limited to inferring upstream regulatory elements using gene sets, but no methods are available for deducing the downstream regulatory elements of transcriptional regulators. There is a pressing need to develop approaches that take into account the bidirectional regulatory relationships of cis-regulatory elements.

The integration of epigenomic multi-omics data is fraught with complexity, including the presence of cross-model and noise, as well as potential non-linear relationships between samples. Existing algorithms only use traditional regression-based methods, such as Lisa, and do not consider the effects of cross-model and noise when integrating multi-omics data. Moreover, the relationships between multi-omics data are not simple linear problems, but a complex network. Deep learning algorithms have achieved great success in solving these specific biological problems^11,12^. The initial step in employing data-driven deep learning approaches involves the extensive collection and processing of epigenetic data. Previously, we developed several epigenetic regulatory databases, such as TcoFBase^13^, CRdb^14^, TFTG^15^, SEdb^16^, and ATACdb^17^, which can extend the scope of epigenomic data. TcoFBase, CRdb, and TFTG, as transcription regulation databases, have collected a large amount of transcription regulator data, while SEdb and ATACdb, as epigenomic databases, have collected the most comprehensive enhancer and chromatin accessibility data. By integrating a large amount of epigenomic resources, we have constructed the most comprehensive epigenomic feature library. Integrating such rich epigenomic data with cutting-edge deep learning techniques presents an unprecedented opportunity to unravel the complex landscapes of the epigenome.

In this study, we proposed a novel data-inspired deep learning framework termed TRAPT (Transcription Regulator Activity Prediction Tool), which can leverage large-scale epigenomic datasets, assimilating advanced knowledge distillation models and graph convolutional neural networks. We have designed a multi-stage fusion deep learning approach to concurrently integrate the signals from upstream regulatory elements within gene sets and the downstream regulatory elements of transcriptional regulators (TRs), in order to obtain an optimal representation of TRs activity and predict key TRs for gene sets with context-specific regulation. To assess the effectiveness of our method, we predicted transcription factors, co-factors and chromatin regulators on up to 570 TR knockdown/knockout datasets from KnockTF^18^ database and 502 TF binding datasets from GTRD. Benchmark tests were conducted against established tools such as Lisa, BART, i-cisTarget and ChEA3, and our results demonstrated that TRAPT outperforms these in predicting TR activity. We also leveraged TRAPT in the study of Alzheimer’s disease, successfully identifying key transcriptional regulators such as REST. We ultimately applied TRAPT to datasets from human cell development and normal human tissues. TRAPT successfully predicted critical regulatory factors controlling cell fate determination as well as tissue-specific regulators.

## Results

### TRAPT overview

TRAPT is a multi-omics integration framework designed for inferring transcriptional regulator activity from a set of query genes. TRAPT employs a multi-stage fusion strategy to address the issues of incomplete cis-regulatory profile coverage and TRBP problems. By leveraging two-stage self-knowledge distillation to extract the activity embedding of regulatory elements, TRAPT can predicts key regulatory factors for sets of query genes through a fusion strategy. The TRAPT framework comprises four main steps: (1) Calculating the epigenomic regulatory potential (Epi-RP) and transcriptional regulator regulatory potential (TR-RP) (**Fig. 1e**); (2) Predicting downstream regulatory element activity of each TR (**Fig. 1a**); (3) Predicting the context-specific upstream regulatory element activity of the queried gene set (**Fig. 1b**); (4) Integrating the predicted regulatory element activity from steps 2 and 3 for predicting the activity of TRs (**Fig. 1c, d**).

**Fig. 1.**
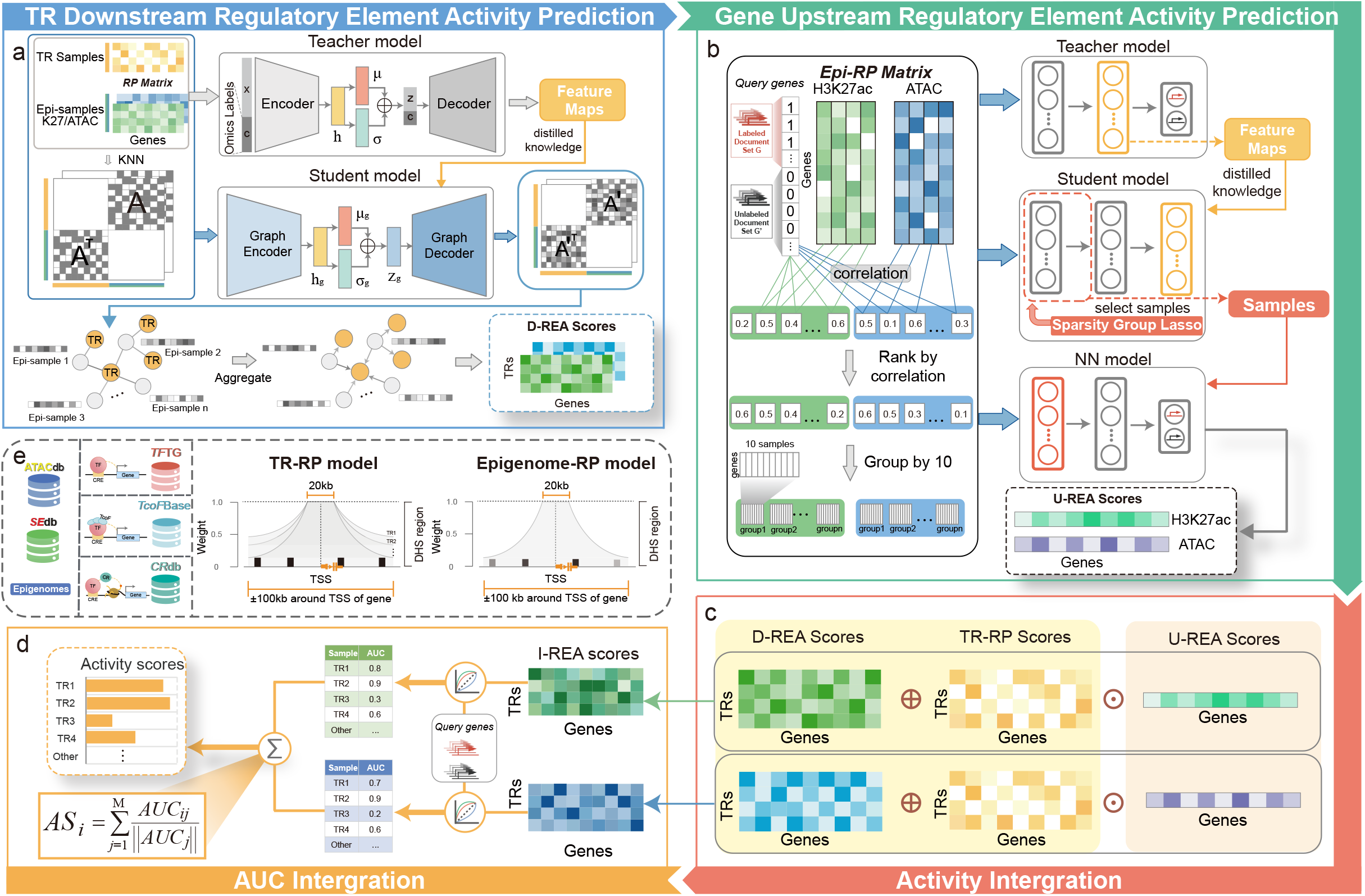
Overview TRAPT. **a** TRAPT predicts downstream regulatory elements activity associated with the TRs. The inputs for TRAPT consist of preprocessed TR-RP matrix and Epi-RP matrix, which are integrated to form an integrated regulatory potential matrix and an adjacency graph. Firstly, we use a conditional variational autoencoder as the teacher network to learn the latent representation h. Then, we employ a graph variational autoencoder as the student network to reconstruct the TR-epigenome adjacency graph by learning its own network structure information and latent feature representation from the teacher network. Finally, we perform an aggregation operation using the reconstructed TR-epigenome adjacency graph and the input Epi-RP matrix to obtain the downstream regulatory element activity matrix. **b** TRAPT predicts upstream regulatory elements activity associated with the queried genes. Firstly, the query gene vector is correlated with each epigenomic sample using Pearson correlation. The samples are grouped based on their correlation values, and the grouped regulatory potential matrix is used as the input for the model. Then, self-KD model is used to select important epigenomic samples. We use a teacher model to extract feature maps, then employ a student model accompanied by SGL constraints to select non-redundant epigenomic samples. Finally, by retraining a nonlinear neural network model with the selected features, we obtain the upstream regulatory element activity matrix. **c** The predicted downstream regulatory element activity matrix, upstream regulatory element activity matrix, and TR regulatory potential matrix are integrated through matrix operations to obtain the I-REA matrix. **d** Firstly, for each TR sample in the I-REA matrix, the AUC score is computed with the query gene set, and the TR AUC scores from all epigenetic groups are integrated to obtain the final activity score for each TR. **e** The epigenomic data is sourced from our previously developed ATACdb and SEdb databases; the TFs, CRs and TcoFs data come from our previously developed TFTG, CRdb and TcoFBase databases. We used the TR-RP regulatory potential model to calculate the TR-RP matrix, and the Epigenome-RP regulatory potential model to calculate the Epi-RP matrix.

To calculate the regulatory potential of epigenomes and TRs, we collected over 20,000 epigenomic sample datasets, including a substantial number of ATAC-seq, H3K27ac ChIP-seq, and TR ChIP-seq datasets, followed by rigorous data preprocessing. Compared to previous methods, TRAPT possesses a more comprehensive and high-quality dataset, with TR data numbering 17,227, which is 2.49 times the amount of Lisa’s dataset and 2.16 times that of BART’s. Furthermore, the chromatin accessibility data we collected and processed exceed the coverage of the largest existing methods by 1.47 times (TRAPT: 1,329; Lisa: 904), and the H3K27ac data is 1.44 times more than that of Lisa (TRAPT: 1,465; Lisa: 1,014) (**Supplementary Fig. 1a, b**). Based on the large-scale epigenome and TR background knowledge library, we calculated the regulatory potential from the epigenome and TR datasets. For epigenomic data, we implemented a uniform weight decay strategy. To effectively differentiate the regulatory scope of various TRs and to provide information on specific regulatory patterns for each TR, we applied a specific weight decay strategy to each TR. (**Fig. 1e**). This effectively differentiates the regulatory scope of different TRs and provides information on specific regulatory patterns for each TR. For a single epigenomic sample, the regulatory potential of a given gene fundamentally represents the aggregated activity of regulatory elements within 100kb of that gene. For a single transcriptional regulator sample, the regulatory potential of a given gene indicates the aggregated activity of transcriptional regulators bound within 100kb of that gene (see Materials and Methods section).

To effectively integrate activities from transcriptional regulators downstream and genes upstream regulatory elements, we divided the activity prediction of regulatory elements into two phases. In the first phase, our goal is to predict the activities of downstream regulatory elements of TRs through integrating Epi-RP and TR-RP. How to effectively aggregate TR-context-specific epigenomic sample signals to predict TR epigenomic activity is a challenge. Meanwhile, methods based on graph convolutional neural networks have demonstrated excellent performance in aggregating neighbor sample information. Therefore, we reformulated the activity prediction problem into a network optimization task. Initially, we use k-nearest neighbours^19^ to construct a heterogeneous network between TRs and epigenomic samples (e.g., CD4+, CD8+ H3K27ac samples) as the initial epigenomic regulatory network (ERN), where the edges of the network represent potential tissue/cell type-specific downstream regulation of TRs. Based on the network, we developed a multi-modal epigenome guided self-knowledge distillation model, named the D-REA model, to optimize the initial ERN. D-REA model can integrate Epi-RP and TR-RP to predict tissue/cell type-specific gene activity of each TR. The teacher model guides the student model to optimize the ERN by extracting low-dimensional embeddings of multi-modal regulatory potential in a self-knowledge distillation manner. Specifically, we input the constructed regulatory potential matrix into a teacher model to learn the joint embedding representations of TR and epigenome samples. We use the teacher model based on a conditional variational autoencoder (CVAE) to effectively integrate regulatory potential features from different modalities. Concurrently, the constructed ERN serves as input for the student model, which learns the low noise cross-modal regulatory potential embedding representations of each TR and epigenomic sample through a variational graph autoencoder (VGAE). During learning, the student model’s parameters are constrained by the omics discrimination knowledge from the teacher model. This enables the student model to reconstructs the ERN as well as amalgamate differential modal features from multiple omics. This approach provides a constrained training environment for the student model, enhancing its resistance to overfitting and generalization performance. The output of the D-REA model is determined by aggregating the activities of regulatory elements in epigenomic samples proximal to the transcriptional regulators. The activity matrix represents the aggregated activity of transcriptional regulators’ downstream regulatory elements near genes, with higher values signifying a more intense level of transcriptional activity in the vicinity of the genes.

In the second phase, our objective is to predict the activities of context-specific upstream regulatory elements of query genes. To accomplish this, an effective strategy is to select significant epigenomic samples from data with noisy and integrate these chosen samples^9,10^. The ultimate integration of Epi-RP signifies the activities of upstream regulatory elements of genes. Therefore, we constructed a self-knowledge distillation model constrained by low-dimensional epigenomic embeddings, named the U-REA model, to infer the activities of upstream regulatory elements of genes. The U-REA model generates low-dimensional nonlinear representations of epigenomic data through a teacher model, while the student model learns these low noise representations and constrains their weights, thereby endowing the U-REA model with the ability to select key epigenomic samples. More specifically, the Epi-RP matrix is used as the model’s input, with a set of query genes and background genes serving as labels for the teacher model. As the model trains, the teacher network is able to extract features of regulatory elements potentially associated with the query genes, and these extracted features are used as softening labels for the student model. The student model learns this low-dimensional epigenomic embedding representation extracted by the teacher model using a network architecture equipped with sparse group lasso (SGL)^20^. By grouping the regulatory potential matrix based on sparse group lasso according to its relevance to the query genes, the student model can impose sparsity constraints both within and between groups. Samples within a group represent highly similar regulatory element profiles, which may contain highly redundant samples. Note that compared to most linear epigenomic sample selection methods^9,10^, the U-REA model employs a non-linear deep learning strategy with accounting for redundant information present in similar epigenomic samples by incorporating a constraint scheme based on sample similarity. This approach allows for more precise selection of non-redundant and non-linearly combined epigenomic samples. Based on the regulatory elements in the selected epigenomic samples, we ultimately constructed another multi-layer neural network model, and the fitted activities of these elements serve as the output of the U-REA model. The output of the final model contains activity vectors that encompass context-dependent information, originating from a specific content-specific gene set. The variation in the values within these vectors represents the specificity of chromatin accessibility (ATAC) and activity (H3K27ac) states associated with the genes’ locations.

By integrating the outputs of the D-REA and U-REA models, we obtain combined regulatory element activities that encapsulate information from both context-free upstream transcriptional regulators and context-specific downstream target genes. We normalized the outputs of both stages’ models. we obtained the integrated regulatory element activity (I-REA) for both modalities by element-wise addition of the normalized TR-RP matrix to the D-REA matrix, followed by an element-wise multiplication with the U-REA vector. Subsequently, for each TR within both models, we quantified its association with the query gene set using the area under the ROC curve (AUC) score^8^. Finally, we merged the activity scores of the corresponding TRs from both modalities to obtain the final combined activity score. In summary, TRAPT integrates downstream regulatory element activity of transcriptional regulators and upstream regulatory element activity of genes to infer the key transcriptional regulators that regulate the queried gene set. We have outlined the overall architecture and concept of the model; specific implementation details can be found in the Materials and Methods section.

### TRAPT demonstrates state-of-the-art performance on benchmark datasets

To comprehensively evaluate the performance of TRAPT, we utilized 570 TR knockdown/knockout datasets from the KnockTF database and 502 TF binding datasets from GTRD for integrated assessments. After quality control, processing, and differential expression analysis, we retained the top-ranked upregulated and downregulated genes of each RNA-seq data as inputs for TRAPT and ultimately evaluated the performance of TRAPT based on the ranking of the target transcription regulatory factors.

We compared TRAPT with several methods that use gene sets as inputs, including Lisa, BART, and i-cisTarget, which utilize TR-ChIP-seq data as a background. Additionally, we evaluated the conventional enrichment analysis method ChEA3, which primarily uses TR-related gene set data as its background. Using various evaluation criteria, such as the number of top 10 TFs recovered, the number of top N TFs recovered, and the overall TF recovery performance, we conducted a comprehensive performance assessment of the models. Based on the metric of the number of top 10 TFs recovered, TRAPT’s performance improved by 13% compared to the second-best method (i.e., Lisa) (**Fig. 2a**). Compared to the classic i-cisTarget method, TRAPT’s performance in predicting the top 10 TFs improved by over 200%. Moreover, compared to conventional enrichment approaches such as ChEA3, TRAPT demonstrates a marked superiority in performance, underscoring the advantages of models predicated on transcription regulator binding. We subsequently calculating the number of correctly predicted transcription factors from cutoff rank 1 to 10 at various thresholds and measuring model performance using the area under the curve (AUC) (**Fig. 2a**). Clearly, TRAPT exhibited the best predictive performance (AUC 0.643). Finally, to evaluate the overall performance of all methods, we calculated the mean reciprocal rank (MRR) scores, a metric used to measure the overall performance of ranking algorithms^21^. The results (**Fig. 2a**) show that TRAPT (MRR 0.067) improved overall model performance by 18% compared to Lisa (MRR 0.057) and by 76% compared to BART (MRR 0.038). Meanwhile, we found that TRAPT significantly outperforms other methods in overall ranking for predicting TRs (**Fig. 2g**). These findings demonstrate TRAPT’s exceptional performance in predicting transcription factors, surpassing previous methods.

**Fig. 2.**
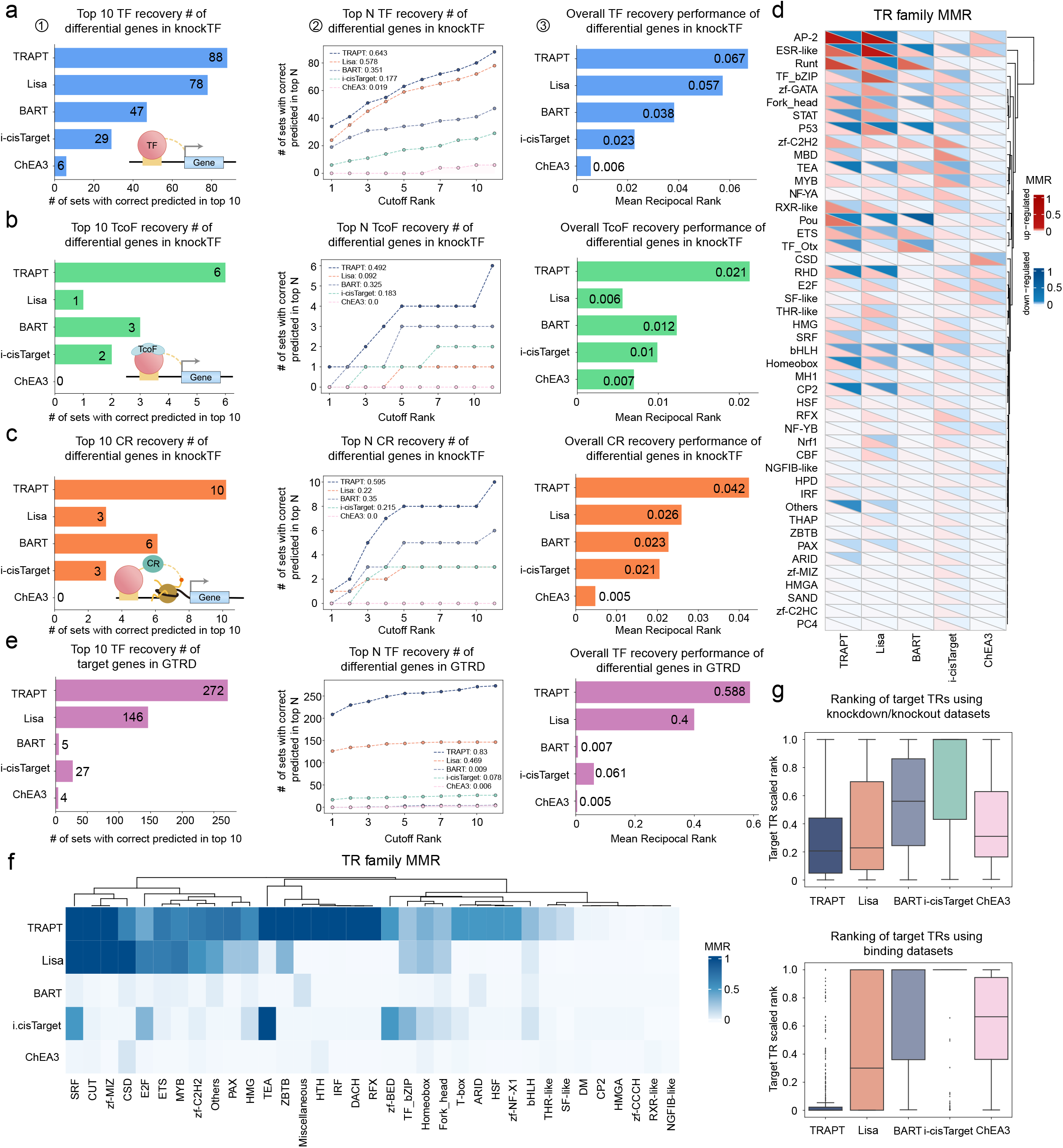
Evaluation of TRAPT and comparative methods on TR knockdown/knockout and TF binding datasets. **a** (1) Shows the number of top 10 TFs accurately identified by five methods: TRAPT, Lisa, BART, i-cisTarget, and ChEA3. (2) Line graph depicting the accurate prediction of TFs from knockdown/knockout experiments across various computational models. (3) Bar graph indicating the MRR scores for TFs, with higher scores reflecting superior performance. **b-c** Subsequent panels maintain the formats of panels (a), extending the analysis to TcoFs and CRs, demonstrating each method’s predictive capability and accuracy. **d** The MRR scores for protein families from TR knockdown/knockout datasets are displayed, with red indicating the upregulated set and blue denoting the downregulated set. The intensity of the color signifies the level of the score. **e** Assess the performance of five methods on TF target genes from the GTRD database, using the same formats as (a). **f** The MRR scores for protein families from TF binding datasets are displayed, where the depth of the color indicates the magnitude of the scores. **g** The box plot illustrates the target TR scaled ranks of across different models in TR knockdown/knockout and TF binding datasets.

Compared to previous methods, which primarily focused on predicting the activity of TFs, our approach, through targeted collection of high-quality ChIP-seq data for TcoFs and CRs (see Materials and Methods section), further predicts the activity of TcoFs and CRs. Although existing methods can predict some TcoFs and CRs, they often conflate these concepts without distinguishing between TFs, TcoFs, and CRs. Our method, however, effectively differentiates among these types of transcriptional regulators. While other methods do not specifically predict non-TF entities, we also benchmarked them and found that TRAPT significantly outperforms existing methods in predicting transcription co-factors and chromatin regulators (**Fig. 2b, c**). We observed a significant decline in Lisa’s performance in predicting TcoFs compared to its second-best performance in TF prediction (**Fig. 2b**), potentially due to Lisa’s collection of extensive data on TFs and CRs but lacking in TcoF data. Moreover, TRAPT’s performance in predicting chromatin regulators far surpasses that of Lisa (**Fig. 2c**). TRAPT’s significant advantages in predicting transcription factors, chromatin regulators and transcription co-factors can be attributed to its use of a multi-stage fusion strategy and its extensive library of transcription regulators and epigenomic backgrounds.

To further test TRAPT’s ability in predicting transcriptional regulators that directly affect genes, we used target genes of 502 TFs from GTRD^24^ (target gene sets bound by transcription factors based on ChIP-seq studies) as inputs for TRAPT, Lisa, BART, i-cisTarget, and ChEA3. The results demonstrated TRAPT’s exceptionally high performance (Fig 2g). We also found that the BART method significantly underperformed on the TF-ChIP-seq target gene datasets compared to its performance on the TF knockdown datasets. Conversely, our method demonstrated enhanced predictive performance on the TF-ChIP-seq target gene datasets, even surpassing the combined scores of four other methods in the top ten performance metrics (TRAPT: 272, other methods total: 182) (**Fig. 2e**). TRAPT also significantly outperformed similar methods in both local and overall performance assessments (**Fig. 2e**).

Subsequently, we explored the performance of different methods across various protein families. The results showed that TRAPT significantly outperformed other methods (**Fig. 2d, f**). Additionally, we found that for certain protein families, TRAPT’s predictive performance completely contrasted between different types of data. For example, the performance on TR knockdown/knockout datasets was notably superior for CP2 and RXR-like compared to TF binding datasets, whereas for families such as zf-C2H2, IRF, THR-like, and CSD, the opposite was true. This indicates the potential for substantial impacts from secondary effects. Finally, a potential issue that may arise from the vast amount of transcription regulators and epigenomic data is the impact on algorithm speed. We have benchmarked the runtime of the TRAP, Lisa, and BART tools (**Supplementary Fig. 1c**). TRAPT’s speed surpasses that of the Lisa and BART algorithms, particularly in predicting the activity of individual transcription regulators (**Supplementary Fig. 1d**).

### Multi-stage fusion strategy boosts transcriptional regulator prediction

To investigate the potential benefits of a multi-stage fusion strategy for predicting transcription regulators, we implemented extensive ablation tests. The U-REA model simulates the activity of upstream regulatory elements for a specific set of input genes, effectively capturing the current epigenetic state of context-specific genes. When the U-REA model was removed from our method, a significant decline in the overall model performance was observed (**Fig. 3a, b and Supplementary Fig. 2a**), showing that the U-REA model reasonably represents the epigenetic state of the input gene set. The D-REA model can predict the epigenomic profile corresponding to the TR, which considers the TR’s preference for the genome under specific conditions. Our approach uniquely considers the activity of downstream regulatory elements associated with TRs. By integrating the binding activity of TRs with the activity of the regulatory elements they bind, our method provides a more comprehensive context-specific insight into TR function. To test usefulness of the model, we removed the D-REA model and observed a significant decline in the overall model performance. This result further demonstrates that considering the specificity of TR binding in regulating element activity is highly effective in enhancing predictive performance. Additionally, by calculating the ratios of transcriptional regulator binding to distal and proximal enhancers, we were able to discern the regulatory preferences of each TR. Consequently, we have developed specific regulatory potential models for each TR to describe their unique regulatory patterns (details in Supplementary Materials). Upon removing the TR-specific regulatory potential model, we observed a decline in the overall performance of the model.

**Fig. 3.**
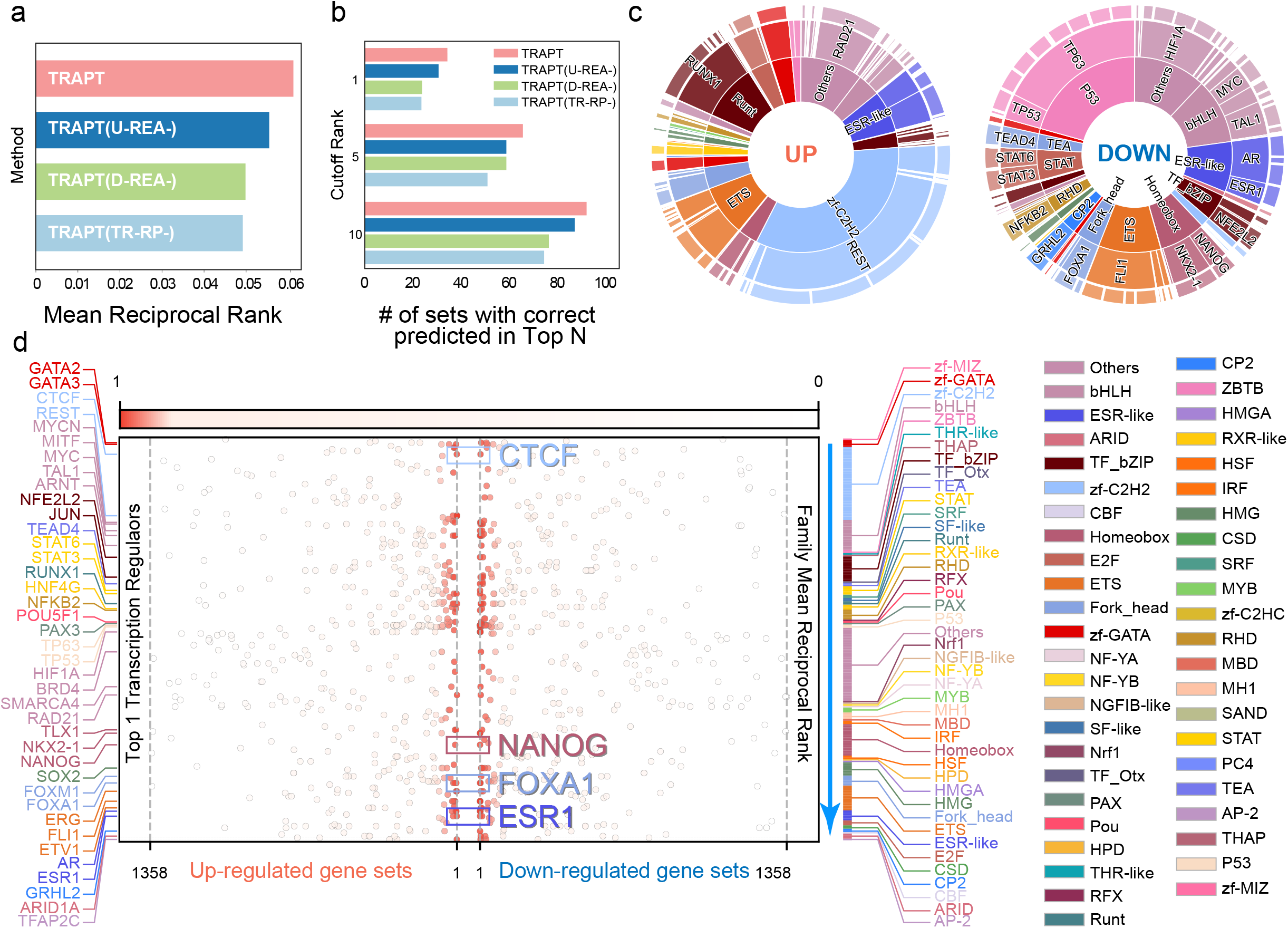
Using the differential gene sets from TR knockdown/knockout experiments by KnockTF, we evaluated the performance of TRAPT. **a** The bar chart represents the MRR scores of the model after gradually removing each submodule. Higher scores indicate better performance for TRs. **b** The grouped bar chart shows the number of correctly predicted top 1, top 5, and top 10 TRs. We progressively removed U-REA, D-REA and the specific TR-RP model to assess the impact of each submodule on the model’s performance. **c** The sunburst chart displays the MRR for all TRs in the upregulated and downregulated gene sets. The top-ranked TR is indicated. The prediction performance is notably better for the downregulated gene set, but the upregulated gene set can also correctly predict transcriptional repressors such as REST. **d** The scatter plot illustrates the ranking of transcriptional regulators in the prediction. The left side represents upregulated gene sets, while the right side represents downregulated gene sets. CTCF, NANOG, FOXA1, and ESR1 have high rankings in both upregulated and downregulated gene sets, indicating their potential dual functions as transcriptional activators and repressors.

We input TR-RP matrix and Epi-RP matrix and generated a ground real network of TR and epigenome using k-nearest neighbours. The D-REA model is designed to optimize the links between TR and the epigenome based on observed links, while also restoring missing links. Therefore, we evaluated the D-REA model from the perspective of link prediction. We divided the observed links into training, validation, and test sets to simulate missing links. By training the model on the training set and then checking the recovery of missing links on the test set, we evaluated the ability of the D-REA model to infer the relationship between TR and the epigenome. We observed that the losses on the validation set of both the teacher and student networks decreased rapidly during the training process (**Supplementary Fig. 2b**), and the area under the receiver operating characteristic curve (auROC) and area under the precision-recall curve (auPRC) values finally reached 0.81 and 0.84 on the test set, respectively (**Supplementary Fig. 2c**). Given the potential for a significant number of false-negative connections in actual scenarios, we masked varying proportions of links to evaluate the stability of the model under different conditions of missing data. The results demonstrated that as the number of masked edges increased (up to 15% maximum), the model’s recovery performance remained robust, with the auPRC exceeding 0.82 and average accuracy (AP) greater than 0.8 (**Supplementary Fig. 2d**). The recovery effect of the model was satisfactory, indicating the stability of the D-REA model in the face of missing disturbances. We subsequently conducted performance testing on the U-REA model. The purpose of the U-REA model is to predict cis-regulatory profiles based on queried gene sets and the Epi-RP matrix. A key challenge is to select crucial epigenomic samples, which are expected to best represent the epigenetic state of the current context-dependent gene set, from redundant data. To tackle this, we computed the model’s performance under different sample selection scenarios. We observed that the rate of performance improvement significantly slowed down after selecting 10 features. This finding aligns with the conclusions of existing research^10^ (**Supplementary Fig. 2e**).

Finally, we compared the performance of TRAPT on upregulated and downregulated gene sets and found that the prediction for downregulated gene sets was better than that for upregulated gene sets (**Fig. 3c**). The result indirectly proved that transcriptional activators are more common than transcriptional repressors^5^ (**Supplementary Fig. 2f, g**). We also found that most transcriptional regulator either act as transcriptional activators or as transcriptional repressors, with a few, such as CTCF, NANOG, FOXA1, and ESR1, having dual functions (**Fig. 3d**). In conclusion, TRAPT can accurately predict transcriptional activators, repressors, and dual functions.

### TRAPT predicts key transcriptional regulators in ESR1 knockdown study

Activating mutations of estrogen receptor alpha (ER/ESR1) are found in approximately 40% of endocrine-resistant ER-positive (ER+) breast cancer cases, making it the most prevalent subtype among breast cancers. As a TF, ESR1 can mediate aberrant expression of a large number of downstream risk genes. To validate the capability of TRAPT in identifying key transcriptional regulators in disease contexts, we applied it to a gene set derived from human MCF7 ER+ breast cancer cells subjected to siRNA-mediated ESR1 knockdown. Upon submitting the differential gene set before and after ESR1 knockdown, TRAPT accurately predicted the transcription factor ESR1 as ranks 1 in the downregulated gene set and 17 in the upregulated gene set (**Fig. 4a**), indicated that ESR1 may possess dual functions of activating and repressing genes in breast cancer^22^. Moreover, TRAPT also identified top-ranked other ESR1 associated cancer transcription factors, transcription co-factors and chromatin regulators such as FOXA1, EP300, and MED1 etc. (**Fig. 4d and Supplementary Fig. 3a**). For example, the transcription factor GATA3 is a key determinant of mammary luminal cell fate^28^. The pioneer factor FOXA1 influences the onset and progression of breast cancer by modulating genomic accessibility^26^. The histone acetyltransferase EP300 acetylates ESR1 enhances the expression of ESR1 target genes in breast cancer cells^27^. Furthermore, the interactions among the top-ranked TRs from the STRING^24^ database showed that they have high-frequency interactions with each other (**Fig. 4b**). The co-expression analysis from the TCGA^25^ breast cancer dataset also reveals a tight relationship among transcriptional regulators (**Fig. 4c**), notably demonstrating a clear co-expression pattern between GATA3, FOXA1 and ESR1. Overall, TRAPT successfully identified ESR1 along with its associated transcriptional cofactors and chromatin regulators. We meticulously explored the potential interactions among these proteins and their genomic binding patterns, demonstrating the efficacy of TRAPT.

**Fig. 4.**
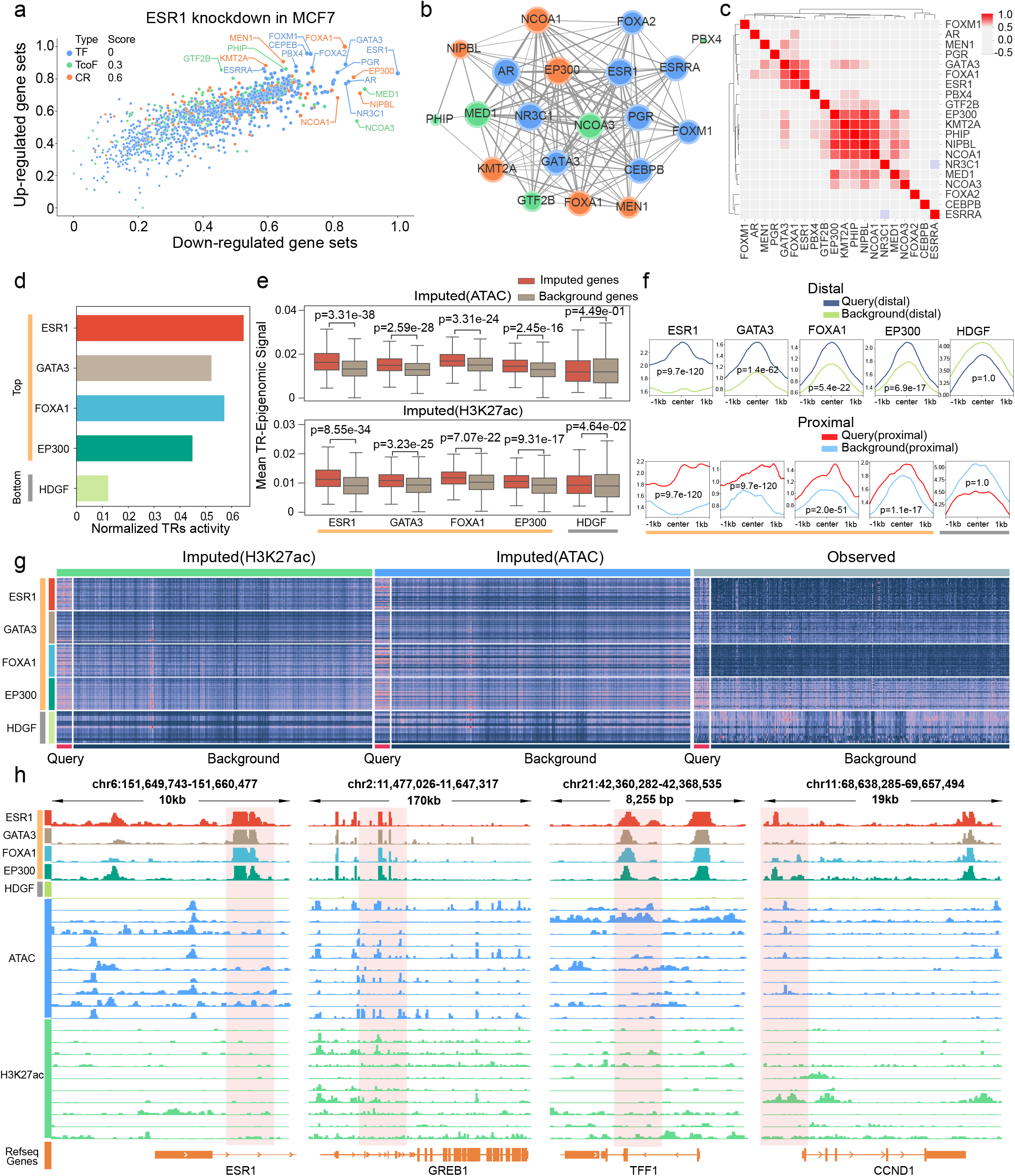
Illustration of the TRAPT framework using downregulated genes from ESR1 gene knockout experiments in gastric cancer and KMCF7 breast cancer. **a** The scatter plot displays the average normalized activity values of 1,358 TRs for upregulated and downregulated gene sets. The size of each data point represents the magnitude of the average normalized activity value, while the colors represent different categories of transcriptional regulators: TFs (blue), TcoFs (green), and CRs (yellow). **b** The network diagram is derived from protein-protein interaction predictions from the STRING database. The size of the nodes represents the degree of the nodes, and the thickness of the edges represents the probability of interaction. **c** The heatmap is derived from the co-expression analysis results of TCGA breast cancer, with the depth of color indicating the degree of correlation. **d** The bar heights represent the current TR normalized activity scores, where ESR1, GATA3, FOXA1, and EP300 are among the top 10 TRs, while HDGF is ranked last. Except for HDGF, with ESR1 having the highest score and HDGF having the lowest score. **e** Comparison of D-REA scores between query genes and background genes revealed significant differences for all TRs, with the exception of HDGF. **f** Aggregated profiles of enhancer marks. Except for HDGF, all TRs marks near the query gene are significantly higher than the background gene. **g** Heatmap of the activity matrix before and after integrating REA scores, demonstrating the differentiation between the query gene and background gene sets. Here, we randomly selected 10,000 genes for visualization. **h** The genome browser displays the tracks of ESR1, GATA3, FOXA1, EP300, and HDGF near the genes ESR1, GREB1, TFF1, and CCND1. We selected the tracks of the top ten epigenomic samples with the highest weights in the reconstructed network for ESR1 and displayed them below, represented by ATAC (in blue) and H3K27ac (in green) tracks.

The D-REA score effectively reflects the epigenetic status of TR, which we combine with its regulatory potential to optimize the activity representation of TR. In theory, representations of transcriptional regulators that incorporate epigenetic information should clearly distinguish the genes they regulate. To validate this, we categorized the identified transcriptional regulators’ D-REA scores into query genes and a background gene set. We found indicate that the top-ranked transcriptional regulators, including the transcription factor ESR1 along with its associated cofactors and chromatin regulators, scored significantly higher in the query gene set compared to the background gene set. ESR1 emerges as the most significant among the transcriptional regulators in both the ATAC and H3K27ac contexts (ATAC: p=3.31e-38, H3K27ac: p=8.55e-34). This indicates that the epigenetic information of ESR1 in cancer can be effectively captured by the representation module of the TRAPT D-REA model. Moreover, we also found that the significance of other top-ranked TRs decreases with their descending rank order, such as NCOA3, NIPBL, and FOXA1. (**Fig. 4e and Supplementary Fig. 3b**). Notably, HDGF was positioned at the bottom in the predictive rankings, and its corresponding significance was substantially reduced compared to other TRs (ATAC: p-value=0.449, H3K27ac: p-value=0.046). These findings underscore the ability of the D-REA scores of transcriptional regulators to effectively discriminate between the genes they regulate. Concurrently, to validate the superior predictive capability of the interpolated I-REA scores over the non-interpolated original regulatory potential scores, we constructed activity profiles for both interpolated and non-interpolated transcriptional regulators. We observed that the top-ranked TRs with high I-REA scores exhibit stronger signals for the corresponding query gene sets (**Fig. 4g**).

ESR1 is capable of binding to enhancer elements that regulate distal target genes, such as ERα-occupied super-enhancers (ERSEs)^25^, while TRAPT adeptly leverages distal information via specialized models of regulatory potential. In order to further investigate and predict the genomic binding characteristics of TRs, we categorized enhancers near the query genes as distal and proximal enhancers and plot the enhancer mark profiles for each predicted transcriptional regulator. We discovered that the predicted upstream transcriptional regulators bind significantly more in enhancer regions near the query genes compared to background enhancer regions (**Fig. 4f**). Conversely, the predicted downstream HDGF binds less in enhancer regions near the query genes than in background enhancer regions. Notably, our analysis demonstrates a strong inclination of GATA3 (Proximal p-value = 9.7e-120 < Distal p-value = 1.4e-62) and FOXA1 (Proximal p-value = 2.0e-51 < Distal p-value = 5.4e-22) to bind proximal to genes, whereas ESR1 and EP300 did not exhibit a comparable preference. Lastly, we visualized the tracks near several significantly downregulated differentially expressed genes for ESR1, GATA3, FOXA1, EP300, and HDGF (**Fig. 4h**). All the predicted upstream transcriptional regulators exhibited conspicuous binding patterns near the genes, and the top ten predicted epigenomic sample tracks highlighted a significant enrichment of regulatory elements near ESR1 binding sites. Additionally, we discerned similar genomic binding patterns for the predicted top transcriptional regulators, while no such pattern was evident for the predicted bottom HDGF (**Fig. 4f**). These findings further substantiate the reliability of TRAPT in predicting transcription factors and their associated transcriptional cofactors and chromatin regulators.

### TRAPT predicts functional transcriptional regulators in post-GWAS analysis of Alzheimer’s disease

Genetic variations at specific DNA positions within transcription factor binding sites can alter the binding affinity or activity of transcription factors, consequently affecting gene expression and cellular processes. Therefore, we applied TRAPT to Alzheimer’s disease (AD) with the aim of identifying key transcriptional regulators impacted by causal variants. To accomplish this, we utilized gene sets associated with AD, as predicted by MAGMA^26^, a tool designed to generate inferred disease gene sets from GWAS summary statistics, as inputs for our algorithm. We then conducted a binding analysis of the disease-associated TRs predicted by TRAPT with the significant causal variants detected by GWAS fine mapping (**Fig. 5a**). Through integrating GWAS data and TR results of TRAPT in AD, we display usefulness of TRAPT in identifying key transcriptional regulators impacted by causal variants.

**Fig. 5.**
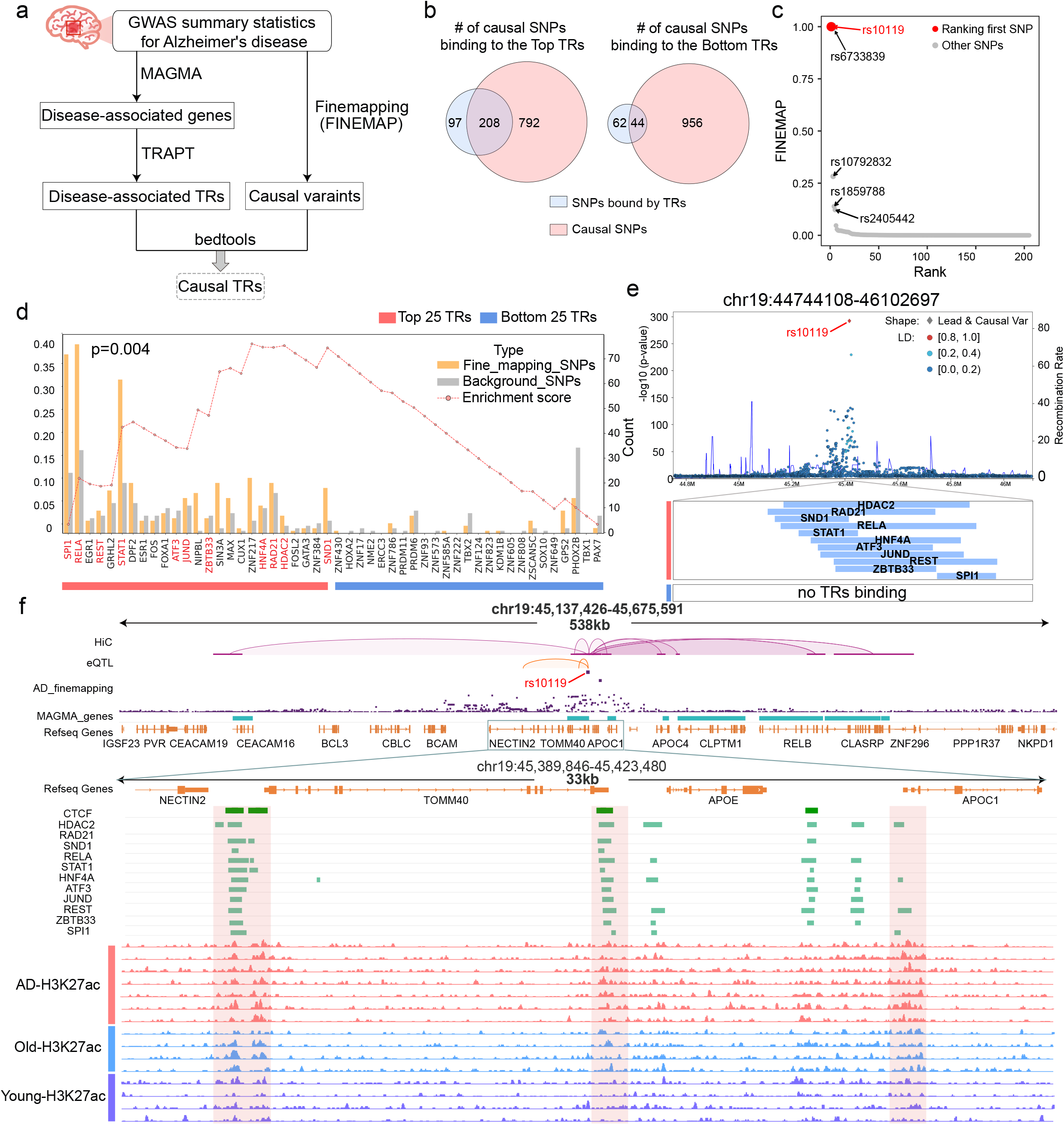
Prediction of functional transcriptional regulators for Alzheimer’s disease using post-GWAS analysis. **a** Alzheimer’s disease analysis workflow. **b** The Venn diagram shows the number of causal SNPs bound near the predicted Top TRs and predicted Tail TRs. **c** The scatter plot displays casual variants bound by top-ranked TRs, with the size of the points indicating the magnitude of FINEMAP scores. **d** The bar chart displays the results from co-localization analysis. The yellow bars represent the number of TRs binding to significant causal variants from fine-mapping, while the gray bars represent the number of TRs binding to background variants. Here, we selected the top 25 and bottom 25 ranked TRs for demonstration. The enrichment line plot represents the binding enrichment of the top 25 and bottom 25 ranked TRs. It can be observed that causal variants tend to co-occur with TRs predicted by TRAPT. **e** Manhattan plot showing the top-ranked causal variant rs10119 obtained from fine-mapping. Additionally, significant gene sets analyzed using the MAGMA software were used as input for TRAPT. The bottom tracks represent transcriptional regulators binding peaks, where EGR1, RELA, REST, and STAT1 are predicted to be ranked in the top 10, while other TRs are predicted to be ranked in the top 25. The bottom 25 TRs in ranking do not show any binding near rs10119. **f** The genome browser displays the chromatin interactions, eQTLs relationships, Top-ranked TRs binding, and the H3K27ac epigenetic landscape from both normal and disease groups.

Specifically, we initially retrieved a GWAS dataset from causaldb^26^, comprising a European population sample of 408,942^27^ as input for fine-mapping. Subsequently, we performed co-localization analysis on disease-associated TRs and predicted causal variants (**Fig. 5a**). Among the 305 SNPs bound by the top 25 predicted TRs of TRAPT, 68.2% belongs to AD-related causal variants (Hypergeometric test p-value=2.48e-12). Conversely, among 106 SNPs bound by the bottom 25 TRs, fewer than half are causal variants (Hypergeometric test p-value=0.971) (**Fig. 5b**). This indicates that TRs ranked highly by TRAPT are more closely associated with AD. Subsequently, to further investigate the relationship between individual TRs and AD, we conducted a more detailed co-localization analysis of each AD-related TR’s binding to causal variants (**Fig. 5c**). The results revealed that the top-ranked TRs such as SPI1, RELA and REST generally had a higher binding affinity to causal variants compared to background variants. For example, SPI1, ranked first by TRAPT predictions, intersected with 71 causal variants, compared to an overlap with only 24 background variants (Hypergeometric test, p=3.6e-4). RELA, ranked second in TRAPT predictions, intersected with 75 causal variants, while overlapping with only 33 background variants (Hypergeometric test, p=2.7e-3) (see Supplementary Table 2). We observed that top-ranking TRs generally exhibit stronger associations with AD-related causal variants. To validate this observation, inspired by the GSEA^28^ algorithm, we developed a new statistical test to verify the reliability of our predicted top-ranked TRs from a statistical standpoint (see Materials and Methods section). Ultimately, we found that the top-ranked TRs were significantly enriched at the top (**Fig. 5d**) (p-value=2e-3), demonstrating that TRs ranked higher were more likely to bind to causal variants compared to those ranked lower.

Next, to identify disease-associated causal variants bound by TRs identified using TRAPT, and to explore their potential associations, we performed co-localization analysis between causal variants and predicted TR binding sites. We retained the overlapping causal variants and ranked them based on FINEMAP scores. Among the selected 1,000 causal variants, 208 are associated with TR binding, with rs10119 ranking as the top variant. (**Fig. 5c**). The functional annotation analysis using VARAdb^29^ revealed that rs10119 is regulated by multiple super-enhancers covering several important genes nearby, such as APOE, TOMM40, and APOC1, and is a risk SNP for Alzheimer’s disease^30^. Subsequently, we conducted a co-localization analysis between rs10119 and the predicted TRs. Notably, in the 1kb region upstream and downstream of rs10119, we observed binding in 9 out of the top-ranked 25 TRs, whereas lower-ranked TRs did not exhibit any binding in these regions. (**Fig. 5e**). A previous study thoroughly validated the effect of REST, a transcription factor, as a universal feature of normal aging in human cortical and hippocampal neurons. It can also potently protect neurons from oxidative stress and amyloid β-protein toxicity. We observed that REST ranks high in TRAPT predictions, and previous studies have already demonstrated that REST plays a crucial role in the development of AD. It inhibits genes that promote cell death and AD pathology, while simultaneously inducing the expression of stress response genes^30^. TRAPT predicts other top-ranked TRs such as SPI1, STAT1, RELA, HDAC2, JUND and HNF4A, which have been previously implicated in the causal relationship with Alzheimer’s disease through prior research ^31–34^. We found that rs10119 is located exactly at a critical position in the chromatin loop structure, with many important TRs predicted by TRAPT binding upstream and downstream. Notably, we observed a substantial number of binding sites from our predicted TRs on TOMM40, APOC1, APOE and CEACAM16 (**Fig. 5f and Supplementary Fig. 4b**). These genes have been proven to significantly influence the onset of Alzheimer’s disease ^35–38^. Meanwhile, by analyzing AD-related H3K27ac ChIP-seq datasets^39^, we discovered that the H3K27ac signal at the rs10119 site is markedly higher in the disease and elderly groups than in the young, disease-free group. This observation suggests that changes in epigenetics could be a key factor influencing the onset of Alzheimer’s disease. Moreover, several transcriptional regulators we predicted are closely associated with epigenetics, such as HDAC2 and ZBTB33 (also known as Kaiso).

### TRAPT identifies transcriptional regulators associated with cell fate and tissue identity

TRs are crucial in coordinating gene expression programs, driving cell fate decisions, and orchestrating intricate biological processes during cellular differentiation and development. The binding affinity of TRs to proximal or distal cis-regulatory elements of downstream marker genes plays a crucial role in maintaining cell identity. To highlight the applicability of TRAPT in cellular development, we employed TRAPT to capture key regulators for marker gene sets of single cell dataset. Briefly, we reprocessed the scRNA-seq data from human hematopoietic stem cell^40^, visualized the first two principal components (**Fig. 6a**), and identified marker genes between different differentiation lineages using the classic model of hematopoietic differentiation landscapes (**Fig. 6b**). We then used TRAPT to identify the top five driving regulatory factors for different cell fate commitment directions (**Fig. 6c**). Notably, among the 42 TRs we identified, 24 were also differentially expressed genes (DEGs). Moreover, we observed multiple TRs appearing across various differentiation lineages, such as EP300, SMAD1, LYL1, SPI1, LMO2, and TAL1 (**Supplementary Fig. 5a**). In addition, some transcriptional regulators were found exclusively in single lineage branches, for example, STAT4 in the LMPs−NK cells lineage branch, a well-known gene regulating intracellular signaling, with STAT4 deletion in NCR1 expressing cells leading to impaired terminal differentiation of NK cells^41^. In the LMPs-pDCs lineage branch, TCF4 is a transcription factor essential for pDC development^42^. We also applied TRAPT to human embryonic stem cells^43^. Following dimensionality reduction and clustering, the cells were categorized into 6 main subgroups (**Supplementary Fig. 5b**). We identified marker genes for each differentiated cluster as well as for the undifferentiated H1 and H9 clusters. These marker genes were then analyzed using TRAPT, resulting in the identification of key transcriptional regulators that cell fate decisions for each differentiated cluster (**Supplementary Fig. 5c**). In the differentiation of H1 into trophoblast-like cells (TB cells), transcriptional regulators such as GATA3, TFAP2A, and GATA2 exhibited higher activity. Notably, GATA2 and GATA3 have been previously validated to be selectively expressed in the trophoblast progenitor cells during early mouse development, where they directly regulate key genes^44^. In the differentiation of H1 into definitive endoderm cells (DE cells), transcriptional regulators like GATA6, SMAD2, and EOMES showed elevated activity. Past studies have revealed that GATA6 works in conjunction with EOMES and SMAD2 to regulate the gene regulatory network associated with human definitive endoderm^45^. TRAPT is capable of effectively identifying driver regulators of cell fate decisions, with the majority of these being cell-lineage-specific transcriptional regulators that have been validated in the literature.

**Fig. 6.**
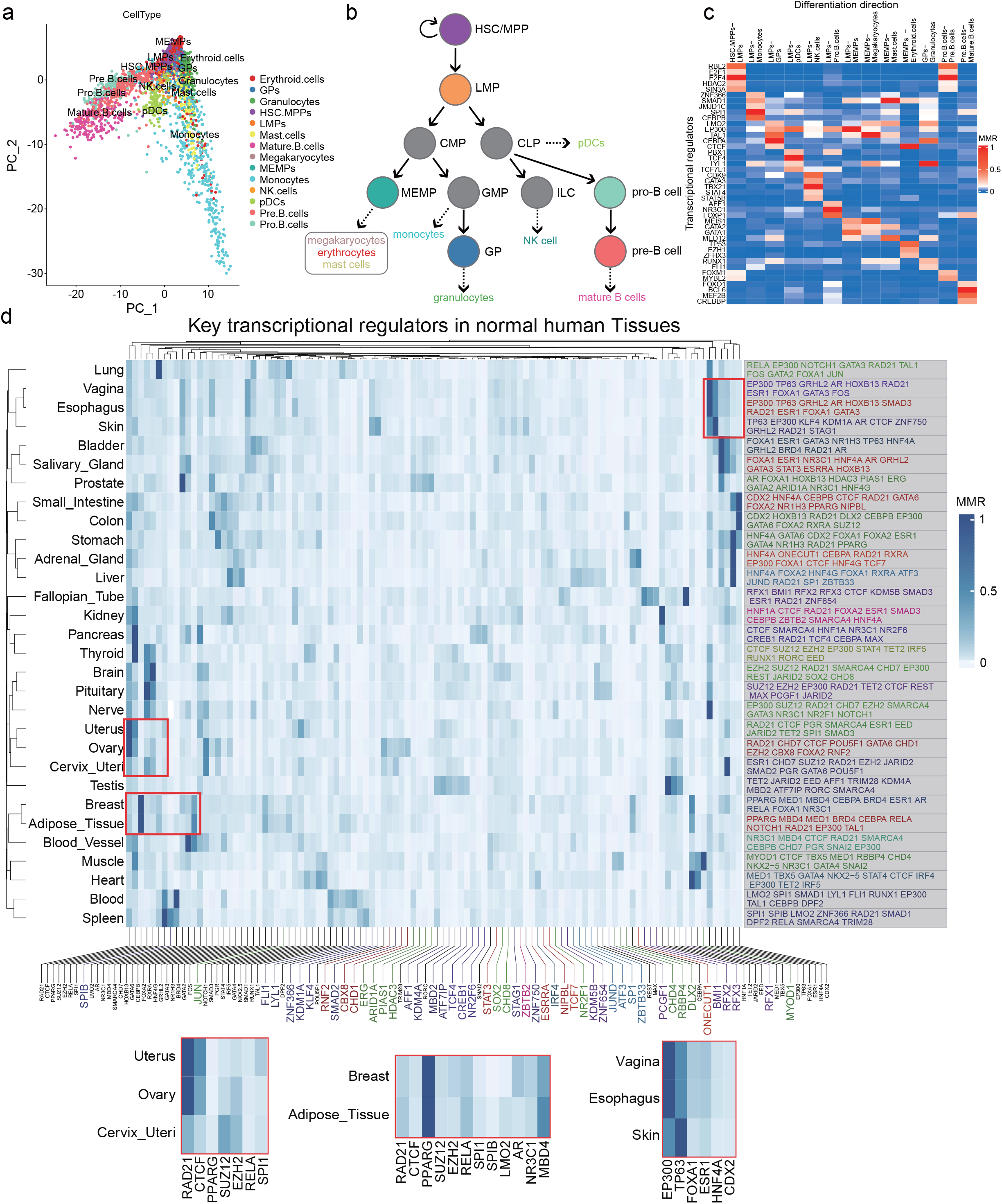
TRAPT identifies transcriptional regulators associated with cell fate and tissue identity. **a** Visualization of principal component analysis (PCA) derived from scRNA-seq data. **b** Classic model of hematopoietic differentiation landscape. **c** Heatmap displays the lineage-specific transcriptional regulator MRR scores of TRAPT across different cell differentiation pathways. **d** The heatmap presents the top 100 TRs by MRR scores, predicted using TRAPT across 30 human tissues. To the right, the top 10 pivotal TRs predicted for each of the 30 tissues by TRAPT are shown. TRs for each tissue are ranked in descending order by their MRR scores, with different colors denoting distinct tissues. Below the heatmap, TRs in varying colors represent tissue-specific TRs from the top 10 predictive TRs. The three smaller heatmaps below highlight the significant TRs within their respective tissues.

Subsequently, we analyzed RNA-seq data from 30 distinct normal human tissues retrieved from GTEx^46^ and utilized limma^47^ to identify the top 500 differentially expressed genes for each tissue, which subsequently were deployed to predict key transcriptional regulators, respectively. Notably, based on the ranks of TRs, most tissue-specific marker regulators were predicted out as expected. For instance, MED1, TBX5, and GATA4 were enriched in heart tissue. MED1 is demonstrated to play an important role in super enhancer formation and activity maintenance. GATA4 broadly co-occupied cardiac super enhancers with TBX5 determine the contractility, calcium handling, and metabolic activity of cardiomyocyte^48^. AR, FOXA1, and HOXB13 have been identified as the top three TRs in prostate, which is consistent with the role of FOXA1 and HOXB13 in regulating normal AR transcription during prostate epithelial development, as well as their involvement in oncogenic AR transcription during prostate carcinogenesis^49^. Furthermore, the results indicated that certain tissues exhibited shared TRs, such as PPARG and CEBPA in breast and adipose tissues^48,49^, TP63 and GRHL2 in skin, esophagus, and vagina tissues^50–52^, suggesting a similarity in the predominant cell types across these tissues. We then integrated the predicted scores of the top 10 TRs from each tissue for hierarchical clustering. Intriguingly, results indicated that TRAPT effectively discerns similarities between tissues (**Fig. 6c**). For instance, breast and adipose tissues formed a cluster due to their predominant composition of adipocytes. Uterus, ovary, and cervix tissues formed a cluster due to their surface and interior being covered by epithelial cells. In addition, we offered a list of predicted top 10 key transcriptional regulators for each tissue (Supplementary Table 4).

In conclusion, TRAPT efficiently predicted key transcriptional regulators in cell fate and across 30 human normal tissues, indicating the capability of TRAPT in processing gene sets yielded from multiple phenotype or conditions data, such as cohort data. A substantial number of these predicted transcriptional regulators have been experimentally validated in the literature for their specific roles in these tissues, further solidifying the reliability of our predictions. TRAPT serves as a potent instrument for exploring and comprehending the functions of key transcriptional regulators in human.

## Discussion

Transcriptional regulators play a crucial role in regulating gene transcription programs, orchestrating the precise timing and spatial distribution of thousands of genes to ensure normal cellular function and development. Importantly, TR-mediated gene programs have distinct epigenetic landscape and are demonstrated as the switch in the changes of cell states and disease phenotypes^1,55^. However, due to the lack of epigenomic data of TRs in many cell types, accurately predicting upstream TRs for given gene sets with biological meanings (i.e., differentially expressed genes or marker genes in single cell studies) remains difficult. To address this issue, we propose a novel deep learning framework named TRAPT that leveraging two-stage self-knowledge distillation to extract the activity embedding of regulatory elements, TRAPT can predicts key TRs for context-dependent gene sets from over 20,000 large-scale epigenomes and TRs background knowledge library. We have demonstrated that TRAPT has improved the accuracy of transcriptional regulator prediction in multiple benchmark datasets. Moreover, TRAPT significantly outperforms the existing methods in overall ranking for predicting TRs such as Lisa, BART, i-cisTarget and ChEA3. We also have successfully identified key transcriptional regulators associated with the disease, genetic variation, cell fate decisions and tissues.

Existing TR prediction methods can be classified into two main categories. The first category is gene set-based methods such as Enrichr, TFEA.ChIP, ChEA3 and MAGIC. These methods use TR-related gene sets as background data and employ statistical tests like hypergeometric distribution to calculate TR significance. However, these methods do not accurately simulate the real binding of TRs and CREs. The second category is based on CREs, which indeed address this issue. They simulate the actual binding scenarios of TRs with CREs near genes to predict TR activity, as seen in methods like i-cisTarget, BART, and Lisa. Nonetheless, these methods still have limitations, primarily due to their neglect of TR binding preferences. TRAPT represents the third category, simultaneously integrating the upstream regulatory elements of gene sets with the downstream regulatory elements of TRs. Based on 1072 TR-related datasets of knockout/silencing/ChIP-seq experiment and multiple evaluation criteria, we found that TRAPT, as the third category method, significantly outperforms other methods in overall ranking for predicting TRs. TRAPT’s significant advantage in predicting transcription factors, chromatin regulators and transcription co-factors can be attributed to its use of multi-stage fusion strategy and its extensive transcription regulator background library. Specially, our method has several advantages and methodological highlights: (1) TRAPT uses a multi-stage fusion to simultaneously address the incomplete cis-regulatory profile coverage and TRBP problems. (2) To mitigate the effects of noisy data, TRAPT employs a feature-based offline knowledge distillation framework^55^ at two stages. During the prediction of D-REA, the teacher network optimizes the student network by extracting embedded representations of cis-regulatory element activities near genes. By grouping the regulatory potential matrix based on sparse group lasso according to its relevance to the query genes, the student model can impose sparsity constraints both within and between groups to treat highly redundant samples. In the prediction phase of upstream regulatory element activity, the teacher network extracts low-dimensional embedded representations of epigenomic information related to the query gene set, guiding the student network in choosing the optimal epigenetic sample set. The KD model is robust to noisy data, significantly enhancing the prediction capability of TR activity. Simultaneously, it maintains the algorithm’s prediction speed even when the amount of TR data is more than double that covered by the existing algorithm with the highest coverage (**Supplementary Fig. 1a, b**). (3) We propose leveraging graph theory to address the challenge of predicting the activity of downstream regulatory elements of transcriptional regulators, which is particularly well-suited for epigenomic datasets with small sample sizes. To investigate the potential benefits of a multi-stage fusion strategy for predicting transcription regulators, we implemented extensive ablation tests. When the U-REA and D-REA model was removed from our method respectively, a significant decline in the overall model performance was observed. We evaluated the D-REA model from the perspective of link prediction. Through validation using test datasets, the D-REA model demonstrated its capability to effectively reconstruct unseen links. Moreover, the D-REA model maintains stable performance even when various proportions of links are masked. These results indicate that the D-REA model can effectively optimize the epigenetic regulatory network.

Through 1072 TR-related datasets, we extensively demonstrated that TRAPT outperforms the current state-of-the-art methods in inferring transcription regulators, especially in the prediction of transcription co-factors and chromatin regulators. We discovered that chromatin regulators, transcription co-factors, and transcription factors exhibit different genomic binding preferences, implying the necessity to consider different transcriptional regulators in transcription regulation studies. In the ESR1 knockout experiment, TRAPT successfully identified ESR1 as top 1 and its associated transcription co-factors and chromatin regulators, such as EP300. We found that the top-ranked transcriptional regulators, including the transcription factor ESR1 along with its associated cofactors and chromatin regulators, scored significantly higher in the query gene set. ESR1, binding distal and proximal enhancers, emerges as the most significant among the transcriptional regulators in both the ATAC and H3K27ac contexts. This indicates that the representation module of the TRAPT D-REA model can capture the epigenetic information of ESR1 in cancer. TRAPT have identified TRs causally related to Alzheimer’s disease near rs10119. Additionally, we observed that TRs ranked higher are more likely to be located near the causal SNPs. We ultimately applied TRAPT to datasets from human hematopoietic stem cells, human embryonic stem cells, and normal human tissues. TRAPT successfully predicted critical regulatory factors controlling cell fate, such as STAT4, TCF4, and GATA, as well as tissue-specific regulators, including MED1, TBX5 and GATA4.

TRAPT offers an enlightening perspective for integrating the epigenetic landscape of transcriptional regulators. However, its performance is still constrained by the quantity of both epigenomic samples. To date, TRAPT encompasses 17,227 datasets related to transcriptional regulators (Supplementary Table 1), and over 3000 sets of H3K27ac ChIP-seq and ATAC-seq samples. This stands as the most comprehensive collection of epigenomic samples to date. However, it does not guarantee that each transcriptional regulator can be paired with its corresponding epigenomic samples. Transcription factors recruit cofactors to perform their functions; the affinity of cofactors can either enhance or reduce the transcription factors’ activity, depending on the co-factor’s role as an activator or inhibitor. Chromatin regulators also influence the activity of transcription factors via chromatin structure modifications. Despite accumulating an extensive amount of data on cofactors and chromatin regulators, we have not yet fully understood the complex effects that arise from interactions between transcriptional regulators, a process that is inherently complex.

In conclusion, TRAPT employs a novel strategy for integrate the epigenetic landscape to predict key transcriptional regulators, and is anticipated to provide instrumental guidance for ensuing research and related computational analysis in the field of transcriptional regulation.

## Materials and methods

### Epigenome and transcriptional regulator datasets and data preprocessing

Gene transcription programs are primarily regulated by the biological activities of transcription regulators and coordinated upstream epigenetic marks, such as histone modifications, and open chromatin states, which can establish and maintain the transcriptional landscape of a cell in response to various internal and external signals. Additionally, studies have demonstrated that epigenetic marks can partially simulate the regulatory shapes of transcription regulators, which can fulfill the gap of transcription regulators coverages. Hence, integrating large scale epigenomic data is beneficial to understand the cell-specific transcription mechanism of genes. In this study, we manually collected and processed ∼20,000 raw epigenomic data of multiple types from numerous data sources, covering more than 1000 tissue/cell types. All these epigenome and transcriptional regulator datasets provide comprehensive regulatory cues to infer gene expression patterns. Detailed processing was described as following contents:

#### H3K27ac ChIP-seq data

The H3K27ac ChIP-seq datasets were obtained from SEdb2.0, which was the previous job of our research group. Briefly, we manually collected 1,739 samples, including experimental and control groups, from NCBI GEO/SRA^57,58^, ENCODE^59^, Roadmap^60^, Genomics of Gene Regulation Project (GGR)^59^ and National Genomics Data Center Genome Sequence Archive (NGDC GSA)^61,62^. We obtained H3K27ac peaks signal data by using Bowtie^63^ and BEDTools^64^ multicov tools to process the raw data.

#### Chromatin accessibility data

The chromatin accessibility datasets were obtained from ATACdb, which was the previous job of our research group. Briefly, we manually collected 2,723 samples to cover multiple tissues or cell types from NCBI GEO/SRA and used Bowtie and BEDTools multicov tools to identify chromatin accessibility peaks signal data.

#### Transcription factors data

The transcription factors ChIP-seq datasets were obtained from TFTG, which was the previous job of our research group. Briefly, we manually collected 11,056 samples, cataloguing a total of 1218 human TFs. Utilizing ChIPseeker^65^ R package and BEDTools, we computed the distribution of various genomic composition and UDHS coverage for each TF.

#### transcription co-factors data

The transcription co-factors ChIP-seq datasets were obtained from TcoFBase, which was the previous job of our research group. Briefly, we manually collected a list of TcoFs in mammals from TcoF-DB v2^66^ and AnimalTFDB 3.0^67^, Meanwhile, we collected 4,246 TcoF-related ChIP-seq datasets in different human cell and tissue types from ReMap, ENCODE, Cistrome^68^, and ChIP-Atlas^69^. We used the liftOver^70^ tool from UCSC to convert all ChIP-seq peak data to the hg38 genome assembly. Utilizing ChIPseeker R package and BEDTools, we computed the distribution of various genomic composition and UDHS coverage for each TcoFs.

#### Chromatin regulators data

The chromatin regulators ChIP-seq datasets were obtained from CRdb, which was the previous job of our research group. Briefly, we processed 2,591 CR-associated ChIP-seq datasets from GEO and ENCODE. We identified the CR binding region using Bowtie, SAMtools^71^, and MACS2^72^, and calculated the distribution of various genomic composition and UDHS coverage for each TcoF using ChIPseeker R package and BEDTools.

Given the large volume of data collected, redundancies originating from the same source may be present. We calculated the peak correlation of all TRs, retaining only one of the samples in cases of a correlation value of 1. Through this filtering process, we ultimately maintained 17,227 unique peak files of transcriptional regulators (Supplementary Table 1).

### TR and epigenome regulatory potential model

The regulatory potential of a gene can be determined by calculating the activity of CREs near the gene^12^. To compute the TR-RP matrix, where rows represent TR samples and columns represent genes, we collected peak data for 17,227 TRs from our earlier-established databases: CRdb, TcoFBase, and TFTG. TRs influence gene expression by binding to CREs located upstream or downstream of the gene. Therefore, we only considered CREs that overlap with the binding sites of TRs for the calculation of gene regulatory potential. The BEDTools was employed to identify regions overlapped with CREs for each TR, which we refer to as potential regulatory elements (PREs). By aggregating the signal values of PREs within 100 kb range upstream and downstream of target genes, we computed the regulatory potential of each gene in each sample, resulting in the TR-RP matrix, where each row corresponds to a TR and each column corresponds to a gene. The regulatory potential of the i-th gene in the j-th sample is defined as:

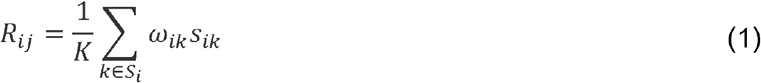

where *ω*_*ik*_ is the regulatory influence of the k-th PRE, located within a 100 kb range of gene j, and *s*_*ik*_ is the signal value of this specific PRE. The weight of each PRE is defined as:

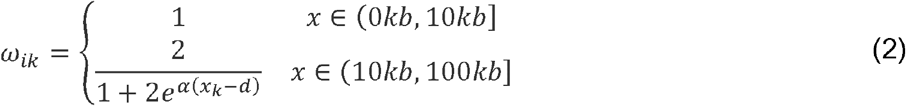

where *x*_*k*_ is the distance between the current PRE and the gene, with the hyperparameter d set to 10 kb.

The parameter *α* controls the decay rate of regulatory influence. The hyperparameters is defined as:

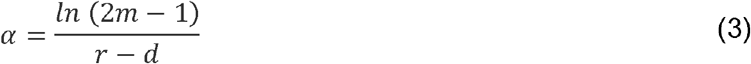

where r is set to 100 kb, and m represents the weight of the regulatory element at distance d. In this case, the weight is determined as the proportion of distal enhancers.

To compute the Epi-RP matrix, where rows represent epigenomic samples and columns represent genes, we used the BAM files from H3K27ac ChIP-seq and ATAC-seq data obtained from SEdb and ATACdb, respectively, and employed the BEDTools multicov tool for counting the reads on the PREs, which produced read signals for all PREs. The computation was performed using the same method, but for each epigenomic sample, we set its m value to 0.01. Furthermore, we used read signals instead of peak signals to compute the gene regulatory potential.

Ultimately, we applied logarithmic standardization on the regulatory potential corresponding to each gene:

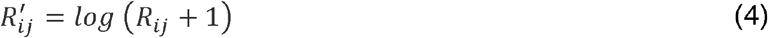

### Predicting the regulatory element activity via upstream transcriptional regulators

In our current module, we leveraged a self-knowledge distillation (KD) model to guide the student model in learning multi-modal epigenomic features and optimizing the epigenomic relationship network. KD originally aimed to compress and accelerate a model by transferring the knowledge from a complex model to a simplified one. In the inference of epigenomic relationship networks, the phenomenon of overfitting frequently occurs. However, recent studies showed that employing self-knowledge distillation significantly enhances the performance of student models and reduces model hallucination issues^57^. Therefore, we propose using KD to infer the downstream regulatory element activity for each TR. Given the distributional differences between TRs and epigenomes, a simple merger is not feasible. To more appropriately extract joint embedding representations of TRs and epigenomes, we utilized conditional variational autoencoders^58^ (CVAEs) as the teacher network. CVAEs have been demonstrated to not only master complex data representations but also outperform at integrating multi-modal data^11^. By incorporating reconstruction error and regularization terms on latent variables during the training phase, CVAEs can learn distinguish feature representations. The model is actualized by minimizing the following loss function:

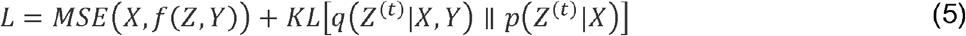

where *X* ∈ *R*^*m*×*n*^ (where m is the number of TR and epigenome samples, and n is the number of genes) represents the feature matrix, assembled by integrating the TR-RP matrix and Epi -RP matrix, *Y* ∈ *R*^*m*×2^ is a one-hot matrix signifies the labels of the two types of omics data, and *KL*[*q* (⋅)║*p* (⋅)] denotes the Kullback-Leibler divergence between the reconstruction network and the (conditional) prior network. Upon inputting the feature matrix and the conditional matrix, we procure a low-dimensional, joint embedding representation *H*^(*t*)^ ∈ *R*^*m*×*h*^ (where h is the dimension of the hidden layer) for TRs and epigenomic samples:

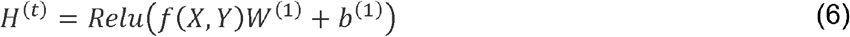

where *W*^(1)^ ∈ *R*^*n*×*h*^ and *b*^(1)^ ∈ *R*^1×*h*^ represent the weights and biases of the encoder’s initial layer, respectively. The complex network presented by the relationship between TR and the epigenome. In order to model this network, we employ VGAE^59^ as the student network and select the nearest 10 epigenomic samples for each TR to construct the adjacency matrix A using k-nearest neighbours^19^, with cosine similarity serving as the distance metric:

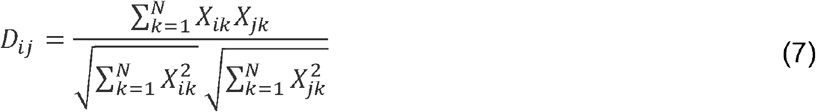

where *D*_*ij*_ ∈ *R*^*m*×*m*^ represents the cosine similarity between the i-th TR and the j-th epigenomic sample, where N equals the number of features (genes). By feeding X and A into the model, we first pass them through the first layer of the GCN encoder to learn the low-dimensional node representations *H*^(*s*)^ ∈ *R*^*m*×*h*^, These node representations not only contain information of a single modality but also encapsulate the relationship information between TR and epigenomic samples. Given that the input is a heterogeneous network, it does not contain any relationship information of a single modality with itself. Subsequently, via the second layer graph convolution, we generate the mean and variance, and ultimately, using the reparameterization trick, we derive the new node feature representation *Z*^(*s*)^ ∈ *R*^*m*×*z*^ (where z is the dimension of the hidden layer in the GCN module). The representation of GCN is expressed as follows:

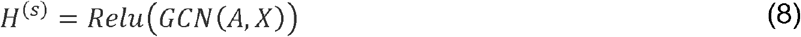

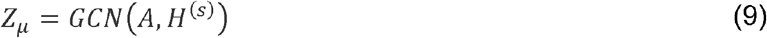

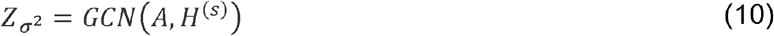

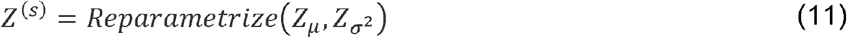

Finally, by utilizing an inner product decoder, VGAE produces a reconstructed adjacency matrix:

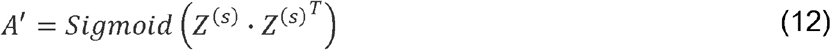

The distillation loss function *L*_*D*_ is defined as:

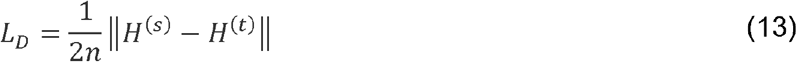

Where ║⋅║ stands for the Euclidean norm. The final loss function L for the student network is defined as:

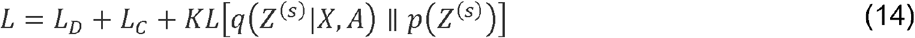

where *L*_*C*_ is the expectation of minimizing the discrepancy between the input and output networks using cross-entropy. *KL*[*q* (⋅) ║ *p*(⋅)] represents the Kullback-Leibler divergence between the reconstruction distribution and the Gaussian prior distribution. For each TR, we predict its corresponding downstream regulatory element activity (D-REA).

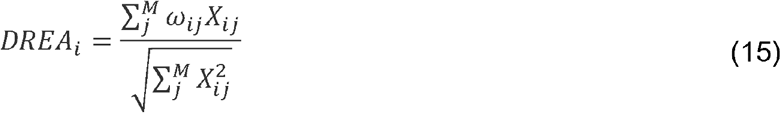

where *X*_*ij*_ denotes the i-th TR corresponding to the j-th neighboring sample, and *ω*_*ij*_ signifies normalized weight of the network edge for the j-th epigenomic sample of the i-th TR. M stands for the number of epigenomic samples. The D-REA matrix represents the aggregated activity of transcriptional regulators’ downstream regulatory elements near genes, with higher values signifying a more intense level of transcriptional activity in the vicinity of the genes.

### Predicting the regulatory element activity via downstream gene sets

There are mainly two methods for predicting upstream regulatory elements of genes. The first method involves using distance to infer regulatory elements near the gene, such as the i-cisTarget approach. The second method employs regression to select epigenomic samples and predict the regulatory element landscape across the whole genome, such as the MARGE method. However, it does not account for the redundancy in epigenomic data and the complex non-linear relationships between samples within the epigenome. Inspired by this and several recent works^60,61^, we propose to use a KD-based strategy to select the most probable epigenomic samples associated with the current query gene set. Initially, we calculate the correlation between the Epi-RP matrix and the query gene set. Subsequently, we ranked the epigenomic samples in descending order based on the magnitude of their correlations. Owing to an accumulation of a large number of epigenomic samples, there may be a plethora of sequencing samples originating from the same tissue. These samples are highly redundant. Consequently, we empirically partition the matrix into sets of 10 samples each. The aim is to group similar epigenomic samples, and enforce sparsity both within and between groups, thus effectively preventing feature redundancy and model overfitting. The grouped Epi-RP matrix is fed into the teacher network. The teacher network is a neural network comprised of three fully connected layers. We predict the query binary gene vector *Z* ∈ *R*^*n*×1^ by using the transposed matrix *X* ∈ *R*^*n*×*d*^ (where n signifies the number of genes and d denotes the number of TR samples) of the Epi-RP matrix. In this process, the query gene set is used as the positive set, and we randomly select 6000 background genes as the negative set. To retain more information, we implement the temperature-scaled sigmoid (TSS) as the activation function in the output layer:

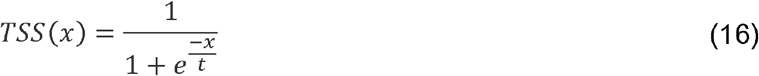

where x denotes the input and t represents the temperature parameter. This function maps the input values to an output value ranging from 0 to 1. As the value of the temperature parameter gravitates towards infinity, the output of the function approximates the output of a standard sigmoid function. Conversely, when the temperature parameter is small, the function’s output alters more gradually within the vicinity of 0 and 1. In this case, we set t=5. The final teacher model is represented as follows:

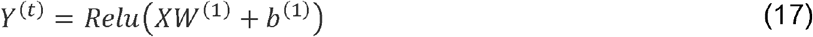

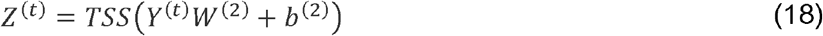

Here, *Y*^(*t*)^ ∈ *R*^*n*×*h*^ (where h denotes the dimension of the hidden layer) signifies the extracted the feature representation of the latent cis-regulatory profiles associated with the query gene set. Between each pair of layers, corresponding weight matrices *W*^(1)^ ∈ *R*^*d*×*h*^, *W*^(2)^ ∈ *R*^*h*×*d*^ and biases *b*^(1)^ ∈ *R*^1×*h*^, *b*^(2)^ ∈ *R*^1×*d*^ exist. We train the student network to predict the low-dimensional feature representation extracted from the intermediate layer of the teacher network by feeding it the same data:

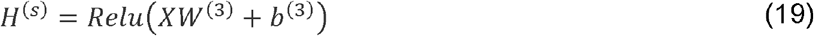

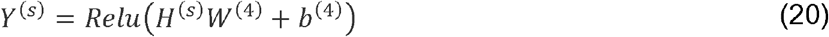

In the first layer of the student network, SGL constraint is used to select important features of cis-regulatory profile by sparsifying the weight matrix through L2 regularization within groups and L1 regularization between groups:

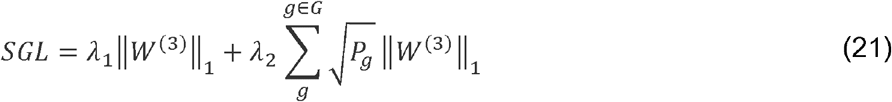

The distillation loss function *L*_*D*_ is defined as follows:

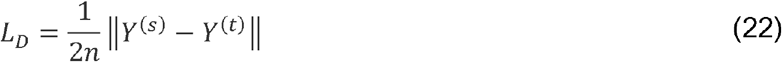

The final loss function L for the student network is defined as follows:

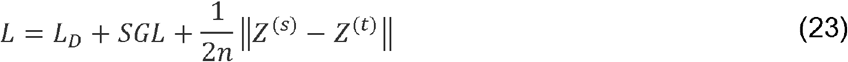

Here, *λ*_1_ and *λ*_2_ represent regularization parameters. We squared and summed the weights of the first layer *W*^(3)^ ∈ *R*^*d*×*h*^of the student network:

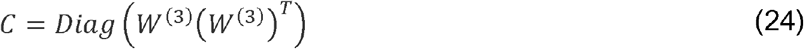

Where C represents the weights of all epigenomic samples, The choice of different sample sizes may impact the prediction results. We selected varying quantities of epigenomic samples based on the weights of the student network model. The aim was to select the most appropriate sample size. We conducted multiple training sessions for the model by selecting varying numbers of epigenomic sample sizes and calculated the auROC to assess the performance of each candidate model. Finally, we found that 10 epigenomic samples were a reasonable number of choices (**Supplementary Fig. 2h**). Utilizing the chosen epigenomic samples, we trained a neural network model, where the input, represented as, *X*′ ∈ *R*^*n*×*d*′^ corresponds to the TR-RP matrix of the selected epigenomic samples, denoted as *d′*. The resulting output is the gene vector *Z*^(*p*)^ ∈ *R*^*n*×1^:

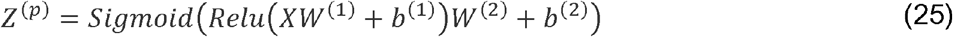

Here we define the loss function as:

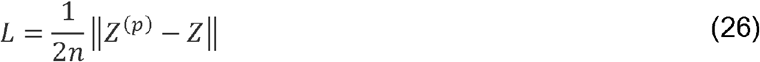

Ultimately, we input X into the previously trained model to derive the predicted gene vector Z, which corresponds to gene upstream regulatory element activity (U-REA). The U-REA vectors encompass context-dependent information, originating from a specific content-specific gene set. The variation in the values within these vectors represents the specificity of chromatin accessibility (ATAC) and activity (H3K27ac) states associated with the genes’ locations.

### Integrating regulatory element activity and predicting transcriptional regulator activity

After obtaining the downstream regulatory element activity and upstream regulatory element activity, we concurrently acquire the downstream regulatory profile information corresponding to TRs and the upstream regulatory profile information corresponding to the queried gene set. Our goal is to obtain the integrated TR regulatory activity that best represents the current gene transcription regulation status. Accordingly, we computed the I-REA for each TR:

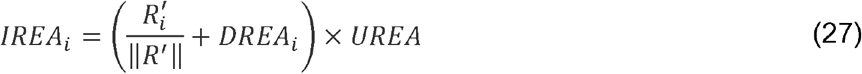

The AUC score has been demonstrated to effectively represent the measurement of transcription factor enrichment^7,8^. Hence, by transforming the query gene set into binary from and computing the AUC for each TR based on its I-REA score, we amalgamated the TRs activity from the H3K27ac and ATAC epigenomes. The resulting activity score was then computed as:

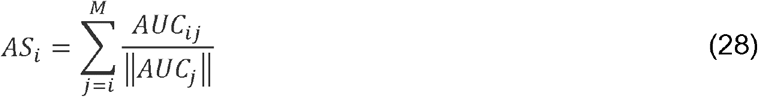

Here, *AS*_*i*_ signifies the final activity of the i-th TR sample, *AUC*_*ij*_ denotes the j-th epigenomic sample AUC score of the i-th TR sample, and M represents the total number of scores.

### Calculate enrichment score and significance in post-GWAS analysis

#### Transcriptional regulator significance

(1) Randomly select 1,000 causal and 1,000 non-causal variants to serve as background variants. Concurrently, select the top 25 and bottom 25 TRs predicted by TRAPT. (2) Utilize the BEDTools intersect tool to compute the number of overlaps between the selected variants and the binding sites of each TR. (3) Calculate the significance p-value of each TR using the hypergeometric test:

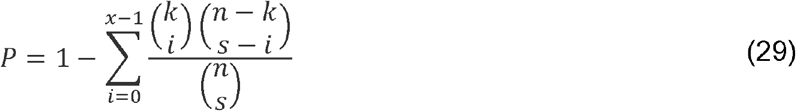

Here, x is the number of causal variants bound by TR, k is the number of variants bound by TR, n is the number of background variants, and s is the number of causal variants in the background.

#### Enrichment score (ES)

(1) Rank the TRs in descending order of activity. (2) ES score calculation:

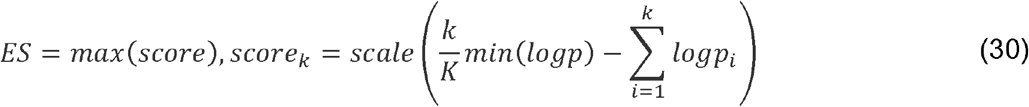

Here, k is the k-th transcriptional regulator, and K is the number of selected TRs. The score is the standard Kolmogorov-Smirnov statistic.

#### Estimating significance

(1) Randomly shuffle the TRs and recalculate the enrichment score as *ES*_*NULL*_ (2) Repeat the shuffle 1000 times and create a corresponding histogram of the enrichment score *ES*_*NULL*_ distribution. (3) Estimate the p-value by calculating the distribution greater than the observed ES.

### Compare TRAPT to similar TR ranking tools

We have collected an extensive array of TR knockdown/knockout datasets from KnockTF, selecting the top 500 upregulated and downregulated differentially expressed genes for analysis. Additionally, we have also curated TF binding datasets from GTRD, retaining all target genes. When comparing TF binding datasets, we removed the data originating from the GTRD in the background TRs ChIP-seq libraries of TRAPT. For the BART algorithm, we utilized the offline toolkit available on their official repository (https://github.com/zanglab/bart2). In a similar vein, for the Lisa algorithm, we employed the offline toolkit accessible on their official repository (https://github.com/qinqian/lisa). For i-cisTarget, we made use of the online analysis tool they provide (https://gbiomed.kuleuven.be/apps/lcb/i-cisTarget/), while for ChEA3, we procured the analysis results via the API online interface available on their official website (https://amp.pharm.mssm.edu/ChEA3).

### Module ablation study

Without disrupting the overall execution of the model, we individually removed the “U-REA model”, “D-REA model”, and “TR-RP model”. We then computed the MRR score for the model after each modification to observe any decline in model performance. The aim was to verify the efficacy of each distinct part. The calculation for the MRR is as follows:

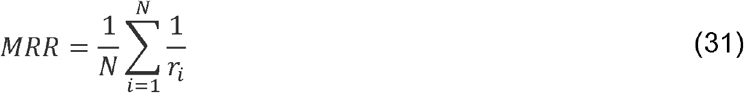

Here, N refers to the number of predicted TRs, and *r*_*i*_ denotes the rank of the current predicted TR.

### Module stability study

Based on the network constructed by k-nearest neighbours, we generated perturbed datasets by masking 2%, 5%, 8%, 10%, 12%, and 15% of the links within the network. These masked links were randomly distributed throughout the entire dataset to simulate real-world scenarios, where we typically cannot determine which interactions are indeed present. During the training phase of the model, we treated these masked positive data as negatives for training purposes. Once the model finished training, we calculated the average precision (AP) to evaluate the model’s predictive performance on the test set. This process helps simulate unknown information in the data, providing a more comprehensive evaluation of the model’s performance. The calculation for the AP is as follows:

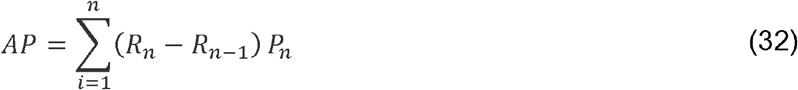

Here, *P*_*n*_ and *R*_*n*_ represent the precision and recall, respectively, sorted by threshold n.

### Software and web tool

TRAPT software was developed using Python 3.11 and has been uploaded to GitHub (https://github.com/LicLab-bio/TRAPT) for user download and utilization. The current iteration of the TRAPT online analysis tool is architected with Python 3.11 and operates on a Linux-based Apache Web server (http://www.apache.org). We employ Django v4.1.3 (https://www.djangoproject.com/) for server-side scripting. The interactive interface is designed and constructed utilizing Bootstrap v4.3.1 (https://getbootstrap.com/) and jQuery v3.2.1 (http://jquery.com). ECharts v5.4 (https://echarts.apache.org/) and DataTables v1.13.2 (https://datatables.net/) are implemented as graphical visualization frameworks, and the sqlite3 lightweight database is deployed for data table storage.

Furthermore, we have developed a corresponding web service (https://bio.liclab.net/TRAPT). The website is designed to accepts gene sets input by users for analysis, allowing easy retrieval of analytical results. We’ve also thoughtfully included an email notification feature. On the results page, the website displays all TR activity scores, as well as the ranking and all individual scores of TRs. Concurrently, the website provides annotation details and relevant quality control information for each transcriptional regulator. Compared to offline tools, online analysis tools offer additional features on the browsing and result analysis pages. The online tools facilitate visualization of the predicted 3D protein structure for each TR, leveraging AlphaFold’s^53^ predictions. Additionally, the online tools incorporate a genome browser^54^ to facilitate user interaction with the genomic tracks associated with each TR.

## Data availability

All the datasets analyzed in this study are publicly available. TR knockdown/knockout datasets from KnockTF and TF binding datasets from GTRD. The protein-protein interaction (PPI) networks from the STRING database (https://string-db.org/). The breast cancer scRNA-seq expression profiles from TCGA (https://portal.gdc.cancer.gov/). The ESR1 knockdown RNA-seq datasets are available in the Gene Expression Omnibus (GEO) repository under accession number GSE37820 (https://www.ncbi.nlm.nih.gov/geo/query/acc.cgi?acc=GSE37820). The GWAS dataset from causaldb (http://www.mulinlab.org/causaldb) and the Alzheimer’s disease-related H3K27ac data are available under GSE65159 (https://www.ncbi.nlm.nih.gov/geo/query/acc.cgi?acc=GSE65159). The human hematopoietic stem cell dataset is available on GitHub (https://gitlab.com/cvejic-group/integrative-scrna-scatac-humanfoetal#data), the human embryonic stem cells dataset is available under GSE75748 (https://www.ncbi.nlm.nih.gov/geo/query/acc.cgi?acc=GSE75748), and the normal human tissues expression profiles from GTEx (https://www.gtexportal.org/home/).

## Code availability

The TRAPT algorithm is implemented in Python. The source code of TRAPT is available at https://github.com/LicLab-bio/TRAPT.

## Acknowledgments

This work was supported by National Natural Science Foundation of China [62171166, 62302206]; Research Foundation of the First Affiliated Hospital of University of South China for Advanced Talents [20210002-1005 USCAT-2021-01]; China Postdoctoral Science Foundation [2019M661311]; Natural Science Foundation of Hunan Province [2023JJ40594, 2023JJ30536]; Clinical Research 4310 Program of the University of South China [No. 20224310NHYCG05].

