## Supplementary Materials for "TRAPT: A multi-stage fused deep learning framework for transcriptional regulators prediction via integrating large-scale epigenomic data"

**Processing and analysis of ESR1 knockdown RNA-seq data**

We downloaded RNA-seq expression profile data from GSE37820 and performed differential expression analysis using the limma R package, retaining the top 500 up-regulated and down-regulated differentially expressed genes for input into TRAPT. We applied Wilcoxon rank-sum test to compute the significance p-value of the integrated D-REA scores for ESR1 and its associated transcriptional regulators in query genes, in comparison to background genes. Leveraging all enhancer sites sourced from SEdb, we categorized enhancers near the query genes as distal (5-100kb upstream or downstream of the gene) and proximal enhancers (within 5kb of the gene). We employed the deeptools^1^ tool to plot the enhancer mark profiles for each predicted transcriptional regulator. Utilizing the Kolmogorov-Smirnov test, we discovered that the predicted upstream transcriptional regulators bind significantly more in enhancer regions near the query genes compared to background enhancer regions.

**Genome-wide association data and methods used in Alzheimer's disease analyses**

Alzheimer's disease GWAS dataset from causaldb. The dataset includes GWAS summary statistics relative to Alzheimer's disease and the corresponding results of statistical fine-mapping. We selected the top 1000 causal variants based on their scores in the FINEMAP^2^ results (Supplementary Table 3), subsequently removing other causal variants from the GWAS results. We randomly chose a same number of variants from the remaining filtered variants as the background variant set. Co-localization analysis is performed using BEDTools.

**scRNA-seq data preprocessing and tissue-specific genes inference**

For the in-depth analyses of the differentiation of human hematopoietic stem cells and embryonic stem cells, the scRNA-seq data were preprocessed using the Harmony^3^ and Seurat package. Batch correction is performed using the ‘RunHarmony’ method from Harmony. The cells with more than 40 zero expressed genes were deleted and the genes expressed on fewer than 3 cells were removed. The top 2000 highly variable genes were selected using the ‘FindVariableFeatures’ method in the Seurat. Finally, the expression values are scaled and centered using the ‘ScaleData’ method in the Seurat. The selection of marker genes is performed using the ‘FindMarkers’ method from Seurat. To obtain tissue-specific genes in normal human tissues, we conducted differential expression analysis on 30 types of tissues from GTEx using the limma^75^ package.

**Differential regulatory patterns of transcriptional regulators**

We performed genomic occupancy analysis of transcription factors, transcription co-factors, and chromatin regulators using the ChIPseeker package, classifying them based on their genomic proportions, particularly in promoter regions (**Supplementary Fig. 6a-c**). Our analysis revealed distinct binding patterns for different types of transcriptional regulators (**Supplementary Fig. 6d**). Transcription factors exhibited a preference for binding near transcription start sites (3' UTR, first exon, and first intron regions). Transcription co-factors showed a predisposition to bind within promoter regions. while chromatin regulators demonstrated a propensity to bind in distal enhancer regions (we use the distal intergenic region as the distal enhancement region). Nevertheless, transcriptional regulators have a substantial proportion of bindings on CREs, such as promoters and enhancers. Further, for transcriptional regulators, their binding on the gene body also constitutes a significant proportion, facilitating directly regulation of gene expression.

Given the differential binding preferences of transcriptional regulators, it is crucial to consider the range of their individual regulatory impact, particularly for chromatin regulators and transcription co-factors, which seem to exhibit potent long-range regulatory capabilities. For example, CTCF largely binds at the boundaries of topologically associating domains (TADs), and its regulatory effect on genes is primarily through long-range regulation. Consequently, in light of the differential regulatory patterns exhibited by each transcriptional regulator, we formulated a transcriptional regulator-specific regulatory potential model. We incorporated the proportion of distal enhancers as the weighting factor for the binding regulatory elements of each transcriptional regulator.

**Transcriptional regulatory elements tend to bind PREs**

DNase I hypersensitive sites (DHS) are structurally open regions on the chromosome that cover a majority of cis-regulatory elements, including enhancers and promoters. To systematically evaluate the binding preferences of TRs in DHS regions, we collected 2,743,279 unique DHS sites from the ENCODE database, which we collectively refer to as potential regulatory elements (PREs). We examined whether the binding regions of TRs from different tissue samples overlapped with these potential regulatory elements and ranked them based on DHS coverage. As expected, the majority of TR binding regions had an overlap with potential regulatory elements exceeding 90% (**Supplementary Fig. 8a**). Additionally, we observed differences in the proportion of DHS coverage for the same TR in different tissues (**Supplementary Fig. 8b**), possibly due to tissue-specific or cell-line-specific epigenetic landscapes that affect TR binding and consequently influence gene expression programs. Given the preferential binding of transcriptional regulators to open chromatin, we used all UDHS sites as background regions rather than the entire genome.

**Transcriptional regulatory patterns**

We classified transcriptional regulation into three main regulatory modes: Direct-Rgulation, Indirect-Regulation, and Effect-Regulation (**Supplementary Fig. 8c**). Direct-Rgulation is defined as TR binding near genes, directly influencing their transcriptional regulation. Indirect-Regulation is defined as TR binding to regulatory elements, indirectly affecting the upstream and downstream genes. Effect-Regulation is defined as the impact of regulatory elements upstream and downstream of a gene, even in the absence of TR binding. By discerning different regulatory modes, we can effectively interpret the synergistic regulatory patterns between transcriptional regulators and regulatory elements by integrating the regulatory potential signals of transcriptional regulators and regulatory elements in the following manner:

$$S=\left\{ \begin{aligned} \begin{matrix} S^{TR} & pattern1 \\ S^{PRE}+S^{TR} & pattern2 \\ S^{PRE} & pattern3 \end{matrix} \end{aligned} \right.$$

Among them, $S^{TR}$ represents the regulatory potential score of the transcriptional regulator, and $S^{PRE}$ represents the regulatory potential score of the potential regulatory element.

**Supplementary Fig. 1 | Evaluation of TRAPT and comparative methods on TR knockdown/knockout and TF binding datasets.** **a** The bar chart illustrates the number of background epigenome samples used by the TRAPT, Lisa and BART methods. **b** The bar chart shows the number of background TR samples used by different methods. **c** The line graph displays the overall runtime of different methods over 50 repeated experiments. **d** The line graph presents the average runtime per TR sample for different methods over 50 repeated experiments.

**Supplementary Fig. 2 | Using the differential gene sets from TR knockdown/knockout experiments by KnockTF, we evaluated the performance of TRAPT.** **a** Grouped bar chart showing the number of correctly predicted genes for the top 1-10 TRs. We progressively removed U-REA (Dark blue), D-REA (green), and specific TR-RP models (light blue) to assess the impact of each submodule on the model's performance. **b** The line graph illustrates the changes in loss on the validation set during the training of the D-REA model. **c** In the test set, the recovery performance is evaluated, with the left graph showing the auROC scores and the right graph displaying the auPRC scores. **d** The line graphs represent the D-REA student model performance at different masking ratios, where the left graph pertains to auROC scores and the right to auPRC scores. **e** The auROC scores corresponding to the scoring models trained with different sample sizes selected by the U-REA student model. **f** Heatmap showing the top 10 predicted TRs in the down-regulated gene set, where brighter colors indicate higher MRR scores. The top-ranked TRs in the down-regulated gene set are predicted to have potential transcriptional activation roles, and TRAPT is also able to predict potential transcriptional co-factors for these down-regulated TRs, such as RAD21 and SMARCA4. **g** Heatmap displaying the top 10 predicted TRs in the up-regulated gene set, which have potential transcriptional repression roles. TRAPT also predicts potential transcriptional co-factors for these TRs, such as ESR1 and FOXA1.

**Supplementary Fig. 3 | Illustration of the TRAPT framework using the example of down-regulated genes in ESR1 gene knockout experiments in gastric cancer and KMCF7 breast cancer. a** The height of the bars represents the current TR MRR scores, where TRs on the left are from the top 10 TRs, and TRs on the right are from the bottom 10 TRs. **b** The comparison of D-REA scores between query genes and background genes indicated significant differences among the top-ranking TRs.

**Supplementary Fig. 4 | Prediction of functional transcriptional regulatory elements for Alzheimer's disease using post-GWAS analysis. a** Manhattan plot showing the fine-mapping rank of the causal variant rs75627662. Additionally, significant gene sets obtained from the MAGMA software were used as input for TRAPT. The top 100 predicted TRs were analyzed for co-localization, and the track below displays TR binding peaks, with GRHL2,DPF2 and FOXA1 being predicted within the top 10. **b** The genome browser displays the chromatin interactions, eQTLs relationships, TRs binding, and the H3K27ac epigenetic landscape from both normal and disease groups.

**Supplementary Fig. 5 | TRAPT identifies transcriptional regulators associated with cell fate and tissue identity. a** The bar chart represents key TRs predicted by TRAPT that are commonly found across multiple differentiation lineages. The height of the bars indicates the frequency of their co-occurrence. **b** Visualization of principal component analysis (PCA) derived from scRNA-seq data. **c** Heatmap displays the lineage-specific transcriptional regulator MRR scores of TRAPT across different cell differentiation pathways.

**Supplementary Fig. 6 | Differential binding patterns of transcriptional regulatory elements across the genome.** **a** Stacked plot displaying the genomic localization analysis of regulatory element binding sites. The percentage of binding sites for TFs is calculated and represented by different colors. TFs are sorted based on the percentage of binding sites near proximal promoters. **b** The binding site percentage of TcoFs was calculated using the same method as employed for (a). **c** The binding site percentage of CRs was calculated using the same method as employed for (a). **d** Box plots illustrating the distribution of binding sites for TFs, TcoFs, and CRs. TFs exhibit a stronger tendency to bind near transcription start sites (3' UTR, first exon, and first intron regions). TcoFs are more likely to bind in the promoter region. As for CRs, they show a preference for binding in distal enhancer regions (intergenic regions).

**Supplementary Fig. 7 | Transcriptional regulatory elements tend to bind PREs.** **a** Bar chart shows the distribution of UDHS coverage for all TRs, with a yellow dashed line indicating 90% coverage. **b** Circular bar chart shows the UDHS coverage of CREBBP, EP300, EZH2, HDAC2, and SMARCA4 transcriptional regulators in different tissues or cell lines. **c** illustrates schematic diagrams of three regulatory modes: (1) Direct regulation. (2) Indirect regulation. (3) Influential regulation.

**Supplementary Table 1:** Details and annotation information for 17,227 transcription regulators.

**Supplementary Table 2:** List of TRs ranked highest and lowest by TRAPT predictions in Alzheimer's disease.

**Supplementary Table 3:** Fine-mapping results.

**Supplementary Table 4:** Top ten ranked transcription regulators identified in various tissues in GTEx.
