## Supplementary figures and images for "TRAPT: A multi-stage fused deep learning framework for transcriptional regulators prediction via integrating large-scale epigenomic data"

### Supplementary Fig. 1

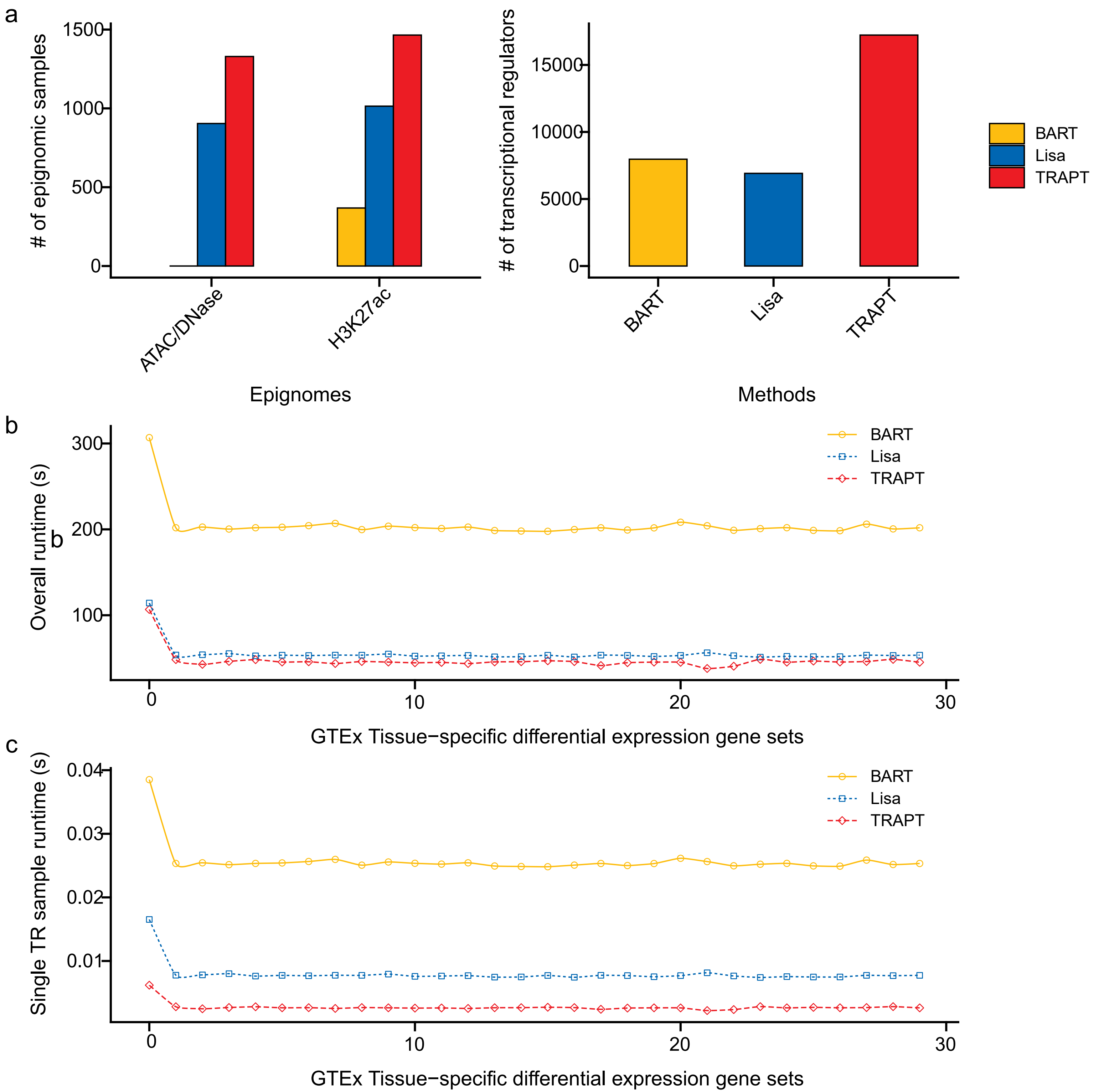

### Supplementary Fig. 2

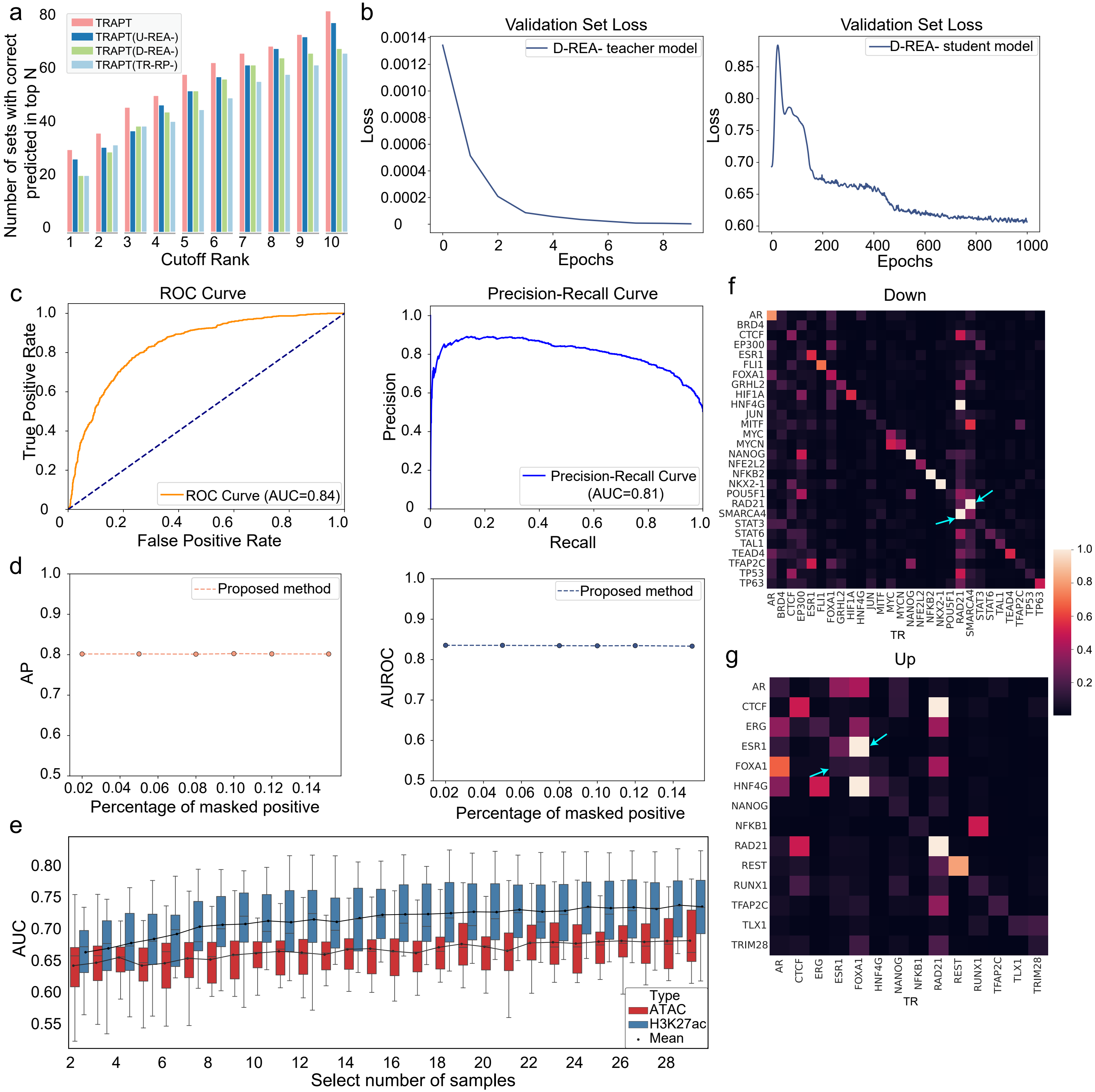

### Supplementary Fig. 3

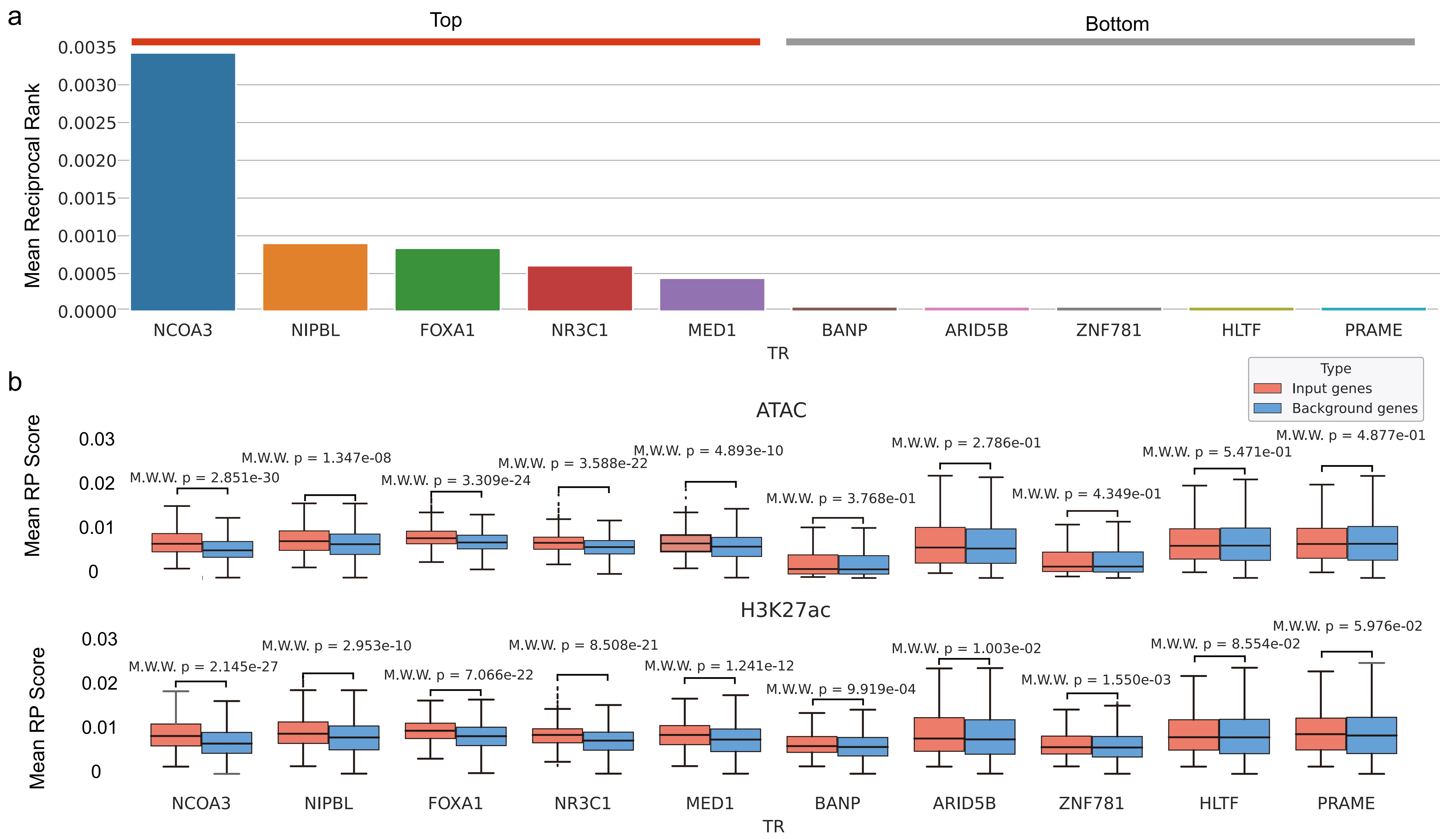

### Supplementary Fig. 4

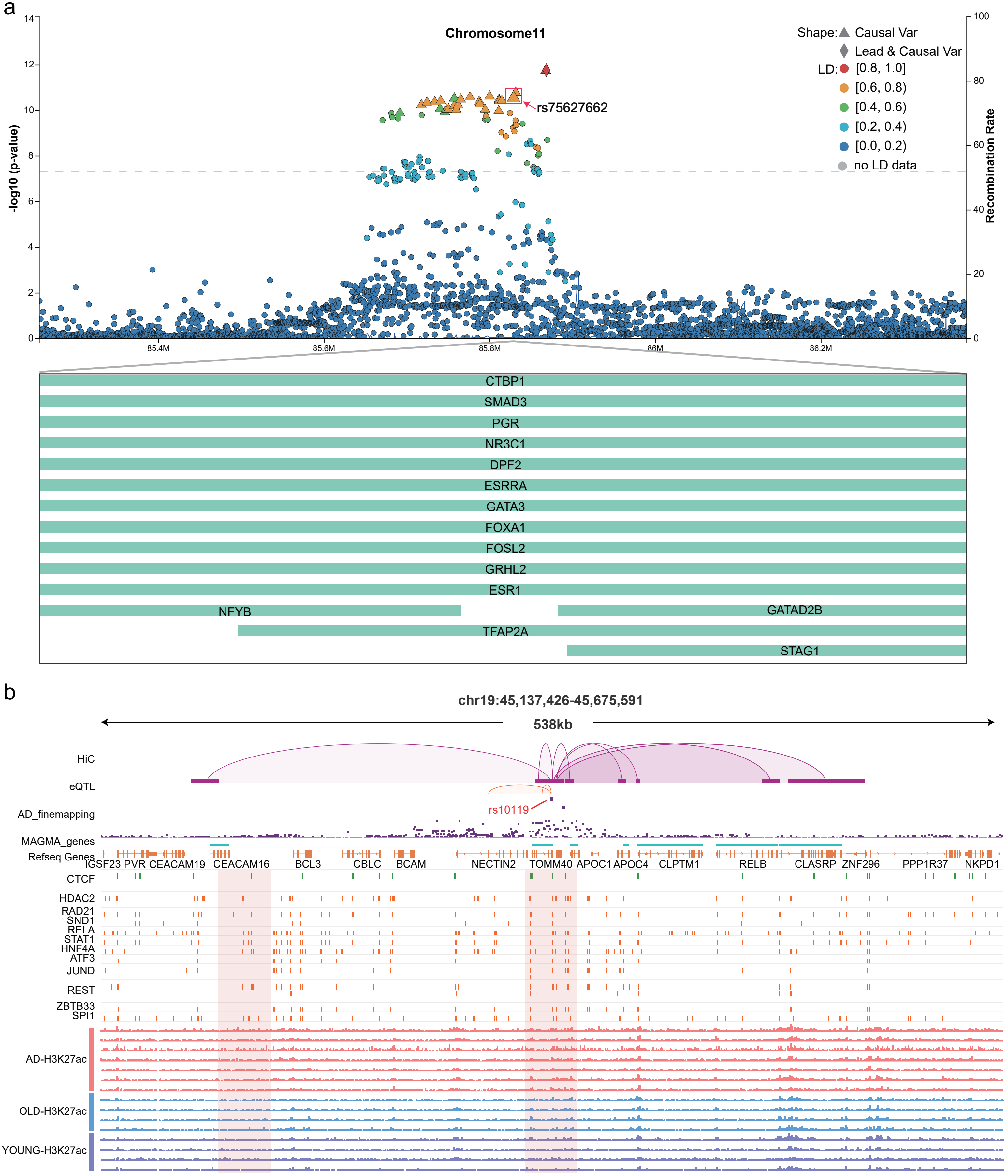

### Supplementary Fig. 5

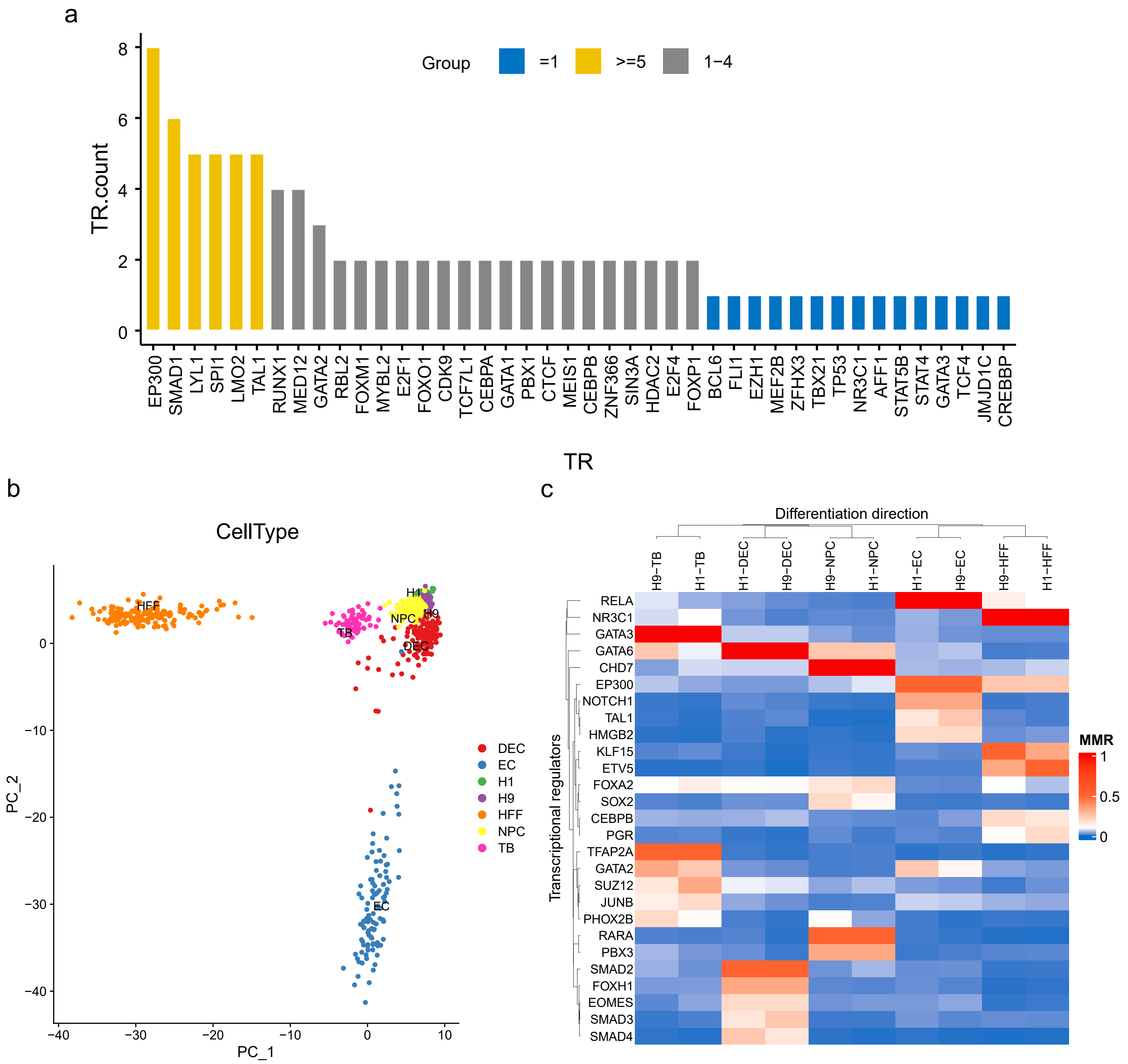

### Supplementary Fig. 6

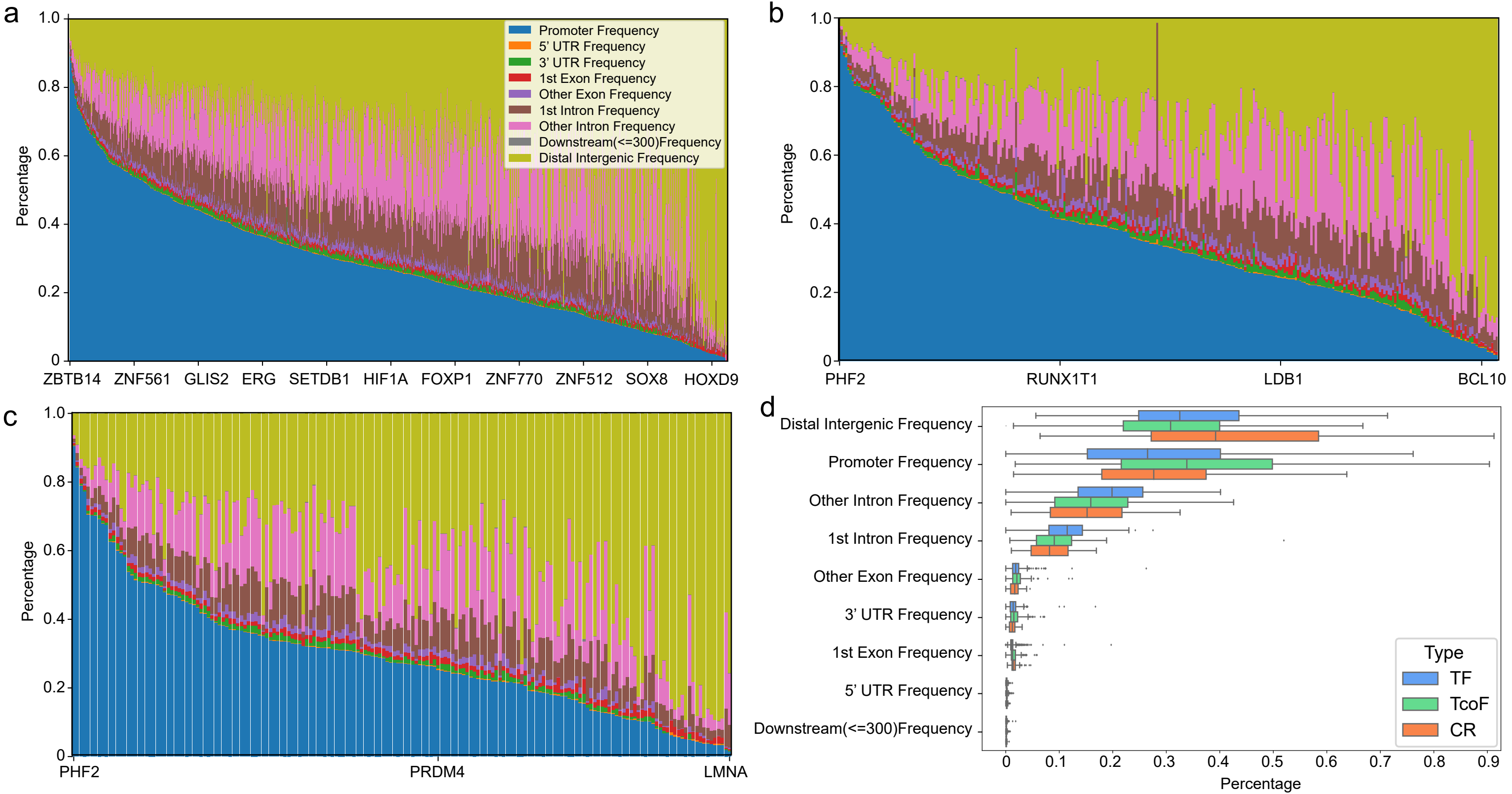

### Supplementary Fig. 7

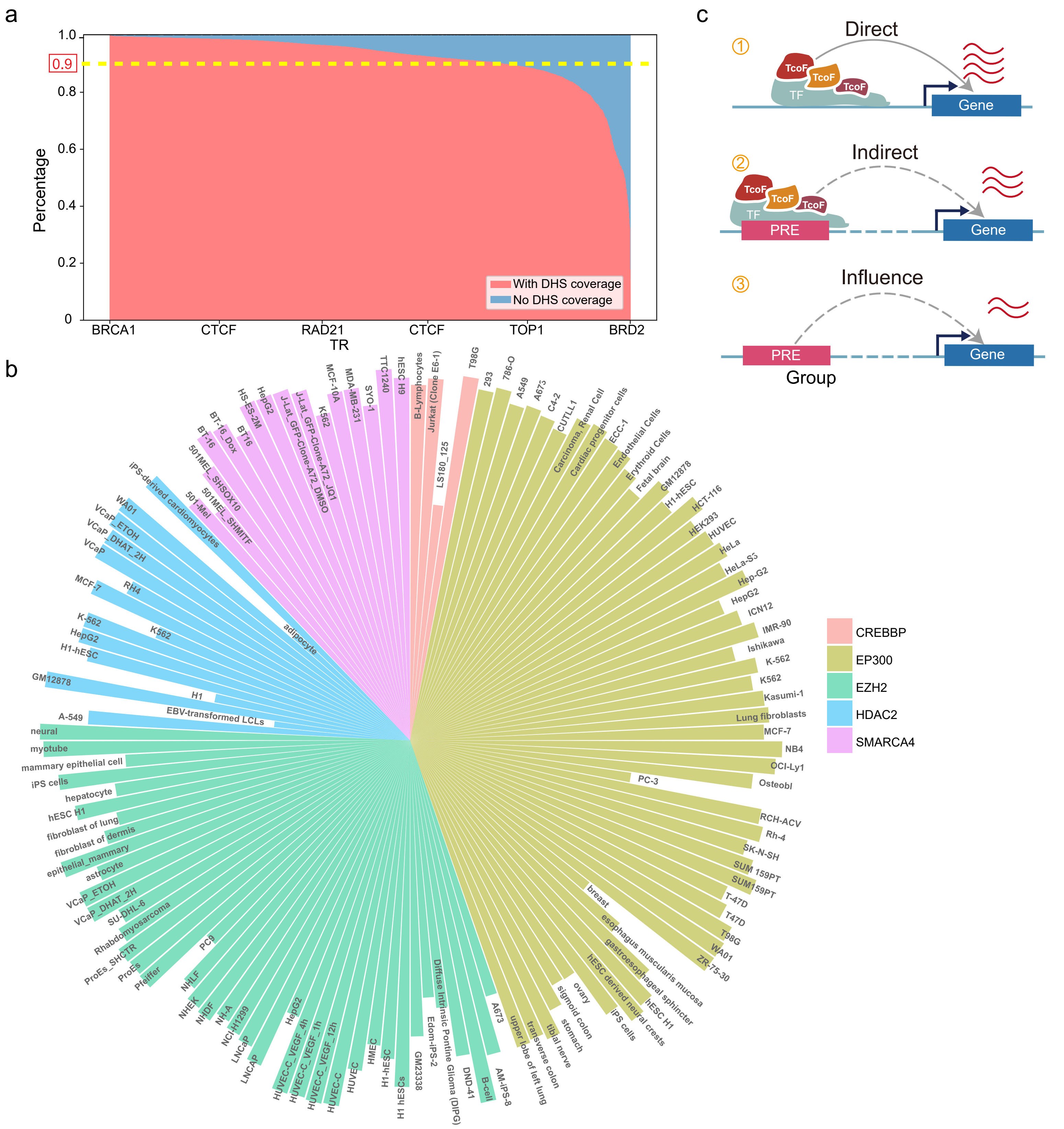
